# Multitrophic interaction networks mediate biodiversity effects on ecosystem multifunctionality

**DOI:** 10.64898/2026.06.15.732252

**Authors:** Georg Albert, Michael Staab, Arong Luo, Perttu Anttonen, Rémy Beugnon, Simone Cesarz, Jingting Chen, Nico Eisenhauer, Alexandra Erfmeier, Felix Fornoff, Pengfei Guo, Werner Härdtle, Lydia Hönig, Lin Jiang, Alexandra-Maria Klein, Yi Li, Yingbin Li, Qi Li, Lingli Liu, Keping Ma, Goddert von Oheimb, Gemma Rutten, Thomas Scholten, Steffen Seitz, Bala Singavarapu, Stefan Trogisch, Ming-Qiang Wang, Pandeng Wang, Donghao Wu, Tesfaye Wubet, Xian Yang, Mingjian Yu, Naili Zhang, Bernhard Schmid, Helge Bruelheide, Xiaojuan Liu, Chao-Dong Zhu, Andreas Schuldt

**Affiliations:** Department of Forest Nature Conservation, University of Göttingen, Göttingen, Germany; Ecological Networks, Technical University of Darmstadt, Darmstadt, Germany; Institute of Ecology, Leuphana University of Lüneburg, Lüneburg, Germany; Key Laboratory of Zoological Systematics and Evolution, Institute of Zoology, Chinese Academy of Sciences, Beijing, China; German Centre for Integrative Biodiversity Research (iDiv) Halle-Jena-Leipzig, Leipzig, Germany; Institute of Biology/Geobotany and Botanical Garden, Martin Luther University Halle-Wittenberg, Halle (Saale), Germany; Research Center for Ecological Change, Organismal and Evolutionary Research Programme, Faculty of Biological and Environmental Sciences, University of Helsinki; Leipzig Institute for Meteorology, Universität Leipzig, Leipzig, Germany; CEFE, Universite de Montpellier, CNRS, EPHE, IRD, Montpellier, France; Institute of Biology, Leipzig University, Leipzig, Germany; College of Biological Sciences, University of Chinese Academy of Sciences, Beijing, China; Institute for Ecosystem Research, Department of Geobotany, Kiel University, Kiel, Germany; Chair of Nature Conservation and Landscape Ecology, Faculty of Environment and Natural Resources, University of Freiburg, Freiburg, Germany; College of Pharmacy, Guizhou University of Traditional Chinese Medicine, Guiyang, China; German Wildlife Foundation, Hamburg, Germany; School of Biological Sciences, Georgia Institute of Technology, Georgia, Atlanta, USA; State Key Laboratory of Vegetation and Environmental Change, Institute of Botany, Chinese Academy of Sciences, Xiangshan, Beijing, China; Key Laboratory of Forest Ecology Management, Institute of Applied Ecology, Chinese Academy of Sciences, Shenyang, China; College of Life Sciences, University of Chinese Academy of Sciences, Beijing, China; Institute of General Ecology and Environmental Protection, TUD Dresden University of Technology, Tharandt, Germany; Institute of Plant Sciences, University of Bern, Bern, Switzerland; Soil Science and Geomorphology, Department of Geosciences, University of Tübingen, Tübingen, Germany; Department of Community Ecology, Helmholtz Centre for Environmental Research - UFZ, Halle (Saale), Germany; Aquatic Geomicrobiology, Institute of Biodiversity, Friedrich Schiller University Jena, Jena, Germany; Key Laboratory of Mountain Ecological Restoration and Bioresource Utilization and Biodiversity Conservation Key Laboratory of Sichuan Province, Chengdu Institute of Biology, Chinese Academy of Sciences, Chengdu, China; State Key Laboratory of Biocontrol, School of Ecology, Sun Yat-Sen University, Guangzhou, China; College of Life Sciences, Zhejiang University, Hangzhou, Zhejiang, China; School of Ecology, Sun Yat-sen University, Guangzhou, China; College of Forestry, Beijing Forestry University, Beijing, China; Department of Geography, University of Zurich, Zurich, Switzerland; Institute of Ecology, College of Urban and Environmental Sciences, Peking University, Beijing, China; Zhejiang Qianjiangyuan Forest Biodiversity National Observation and Research Station, Beijing, China; State Key Laboratory of Integrated Pest Management, Institute of Zoology, Chinese Academy of Sciences, Beijing, China

## Abstract

Biodiversity loss threatens the multifunctionality of ecosystems on which human well-being ultimately depends. Changes in multitrophic species interactions may be key to explaining the ecological consequences of biodiversity loss, but research explicitly linking species interactions and ecosystem multifunctionality remains rare. To assess interaction-mediated biodiversity effects and underlying mechanisms, characterizing the structure of species interaction networks is invaluable. Using comprehensive species interaction and ecosystem functioning data from a large-scale tree biodiversity experiment, we find consistent effects of the structure of species interaction networks on ecosystem multifunctionality across multiple types of antagonistic and mutualistic interactions. While positive effects of network size align with expected positive effects of multitrophic species diversity, positive effects of niche overlap among interacting species and negative effects of highly connected species (i.e. high linkage density) reveal additional, interaction-mediated drivers of multifunctionality. Specifically, the effects of niche overlap suggest benefits of functionally similar species, and the effects of linkage density underscore the importance of specialized interactions in promoting ecosystem multifunctionality. These findings emphasize that to effectively safeguard ecosystem service provisioning, ecosystem management and biodiversity conservation not only need to account for biodiversity changes at multiple trophic levels, but also explicitly for how species interact among each other.

## Introduction

Life on Earth is characterized by an astonishing diversity of species that interact with and depend on each other. As species diversity changes due to human alterations of ecosystems, species interactions within ecological communities change as well^1,2^. These alterations cascade across trophic levels and throughout food webs^1,3–7^. As a consequence, critical ecosystem functions (i.e. ecosystem process rates such as productivity, herbivory, and nutrient cycling)^8^, many of which directly result from species interactions^9^, and essential ecosystem services as the foundation of human well-being^10–13^ are at risk. Biodiversity-mediated shifts in how species interact with each other may thus be a main mechanism behind the often-observed positive effects of species diversity on ecosystem functioning^5,14–18^. However, while many studies have documented general effects of species loss across trophic levels on multiple ecosystem functions (i.e. ecosystem multifunctionality)^5,19–22^, insights into the role that species interactions play in this process are limited to a few trophic levels and ecosystem functions^2,23–26^. At a time when species diversity changes at unprecedented rates^27,28^, the incomplete understanding of the driving forces behind biodiversity-mediated alterations to ecosystem functioning limits our ability to accurately predict the consequences of global biodiversity change and to take evidence-based conservation and restoration action.

Theoretical and experimental studies show that the structure of species interaction networks has major effects on multiple ecosystem functions^7,29^. This is because network structure determines the allocation and transmission of energy and biomass throughout interaction networks^5,9^. Simultaneously, network structure affects long-term ecosystem functioning^30^. As network structure is shaped by species diversity^18,31^ and determines ecosystem functioning^7,29^, it emerges as a key mediator of biodiversity effects on ecosystem functioning. To empirically establish how this mediation takes effect, three aspects of network structure should be considered. First, it is necessary to identify how biodiversity mechanisms (i.e. ecological mechanisms that determine effects of biodiversity on ecosystem functioning) are represented in network structure. Previous biodiversity–ecosystem functioning (BEF) research has particularly highlighted the key role of plant diversity in modulating ecosystem functions^5,15^. Experimental manipulations of plant diversity uncovered the general importance of species complementarity in supporting ecosystem functioning, often attributed to a reduced niche overlap in resource-use among interacting primary producers^32^. Similarly, trophic complementarity (i.e. complementarity in species interactions) alters multiple ecosystem functions in theoretical models, showing that niche overlap in species interactions modifies BEF relationships^17,33^. While niche overlap can stabilize ecosystem functions under changing conditions (insurance effects), it is likely to increase competition and reduce complementarity between species under more stable conditions, thus attenuating biodiversity effects on ecosystem functioning^34,35^. Because complementarity is associated with even contributions of species to ecosystem functioning, but dominant species may increase ecosystem functioning as well^14^, it can be important to also take into account the evenness distribution of species interactions (i.e. interaction evenness).

Second, in addition to biodiversity mechanisms (i.e. trophic complementarity and the dominance of interactions), network structure comprises several dimensions of biodiversity that can affect ecosystem functioning. Theoretical studies have shown that network size (i.e. the number of interacting species in a network) has major effects on multiple ecosystem functions^15,29,36^. Many characteristics of network structure scale with network size, making it an important co-variate to disentangle effects of network structure from effects of network size^31^. Network size can also capture the multitrophic diversity of an ecosystem, which has been shown to be closely related to ecosystem multifunctionality^19,37^. Moreover, not only the species diversity of networks, but also the diversity of each species’ interaction partners (i.e. linkage density) affects ecosystem functioning. While the low trophic niche breadth (i.e. higher linkage density) of specialized interactions may be more effective per capita, the generalism associated with an increased trophic niche breadth (i.e. higher linkage density) can benefit individual species (e.g. flexibility in prey choice) and thus enhance ecosystem functioning^38,39^.

Third, fundamental differences between antagonistic (i.e. one species benefits at the cost of another species, for example during predation) and mutualistic (i.e. species have mutual benefits, for example during pollination) interactions can modify effects on ecosystem functioning^40^. For example, in grassland ecosystems, antagonistic interaction networks are more connected (i.e. higher linkage density) but less nested (i.e. smaller niche overlap)^41^, hence may mediate biodiversity effects differently. Differences between interaction types are especially relevant for ecosystem multifunctionality, as functions associated with antagonistic interactions capture trade-offs (e.g. plant productivity decreasing due to higher herbivory), whereas those associated with mutualistic interactions are more tightly correlated (e.g. reproductive success of plants increases with pollination). Hence, in addition to niche overlap, interaction evenness, linkage density, and network size, the interaction type could alter how species interactions modify biodiversity effects on ecosystem multifunctionality.

Here, we synthesize data on ecological networks and ecosystem functions collected in a large-scale forest biodiversity experiment (BEF-China)^42^ in subtropical China to identify how multitrophic species interactions affect ecosystem multifunctionality and mediate the observed positive effects of species diversity^5,19–22^. Specifically, to assess the simultaneous performance of multiple ecosystem functions we combine 34 measures of ecosystem functions into an ecosystem multifunctionality index that uses the Hill number framework and accounts for correlations between functions^43^. The integrated functions include primary productivity, nutrient cycling, and activity rates of animal and fungal species (Fig. 1a, Supplementary Table 1). Combining ecosystem functions in a multifunctionality index enables us to also account for trade-offs between functions and effects of species interactions that cascade to functions via indirect pathways through higher-order interactions (e.g. predator–prey interactions altering nutrient dynamics^44,45^). In total, we analyzed the network structure (network size, niche overlap, interaction evenness, and linkage density; Fig. 1b, Supplementary Table 2) of 374 interaction networks spanning 11 types of networks, each composed of two adjacent trophic levels (i.e. bipartite networks), with many being based on observed interactions. The networks cover a wide variety of species interactions, both antagonistic and mutualistic, including above- and belowground species, from microbes to predatory arthropods (Fig. 1a, Supplementary Table 3). Thus, our networks represent a substantial proportion of the species pool and their interactions at our study site, allowing for a comprehensive assessment of the role of species interactions in driving biodiversity–ecosystem functioning relationships.

**Fig. 1:**
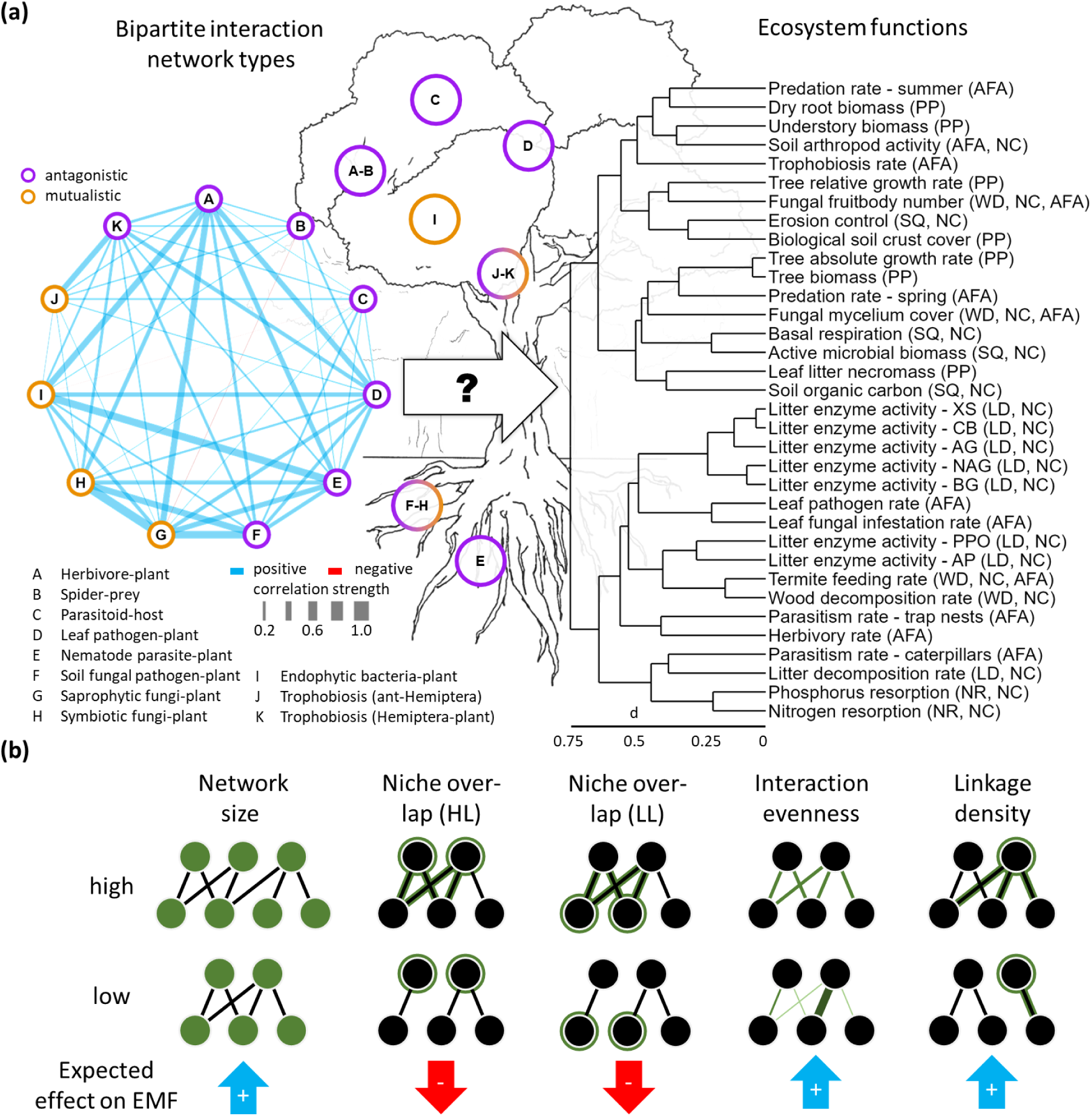
Overview of species interaction networks, ecosystem functions, and network indices. (a) Left graph illustrates correlations among network indices of 11 types of species interaction networks (A-K), averaged across all network indices considered (i.e. network size, niche overlap of higher (HL) and lower trophic level (LL), interaction evenness, and linkage density of bipartite interaction networks; see Supplementary Fig. 1 for individual network indices and Supplementary Table 2 for details on their calculation). Antagonistic (i.e. higher trophic level benefits at the cost of lower trophic level) and mutualistic (i.e. interactions benefit both trophic levels) interaction networks shown as purple and yellow nodes, respectively. Positive correlations are indicated in blue, negative correlations in red. Line width scales with the strength of a correlation. Only correlations with p < 0.05 are shown. Right graph in (a) shows dendrogram of ecosystem functions based on correlation-based distances d used to calculate ecosystem multifunctionality. Ecosystem functions include measures of primary production (PP), nutrient cycling (NC), nutrient resorption (NR), soil quality (SQ), litter decomposition (LD), wood decomposition (WD), and animal and fungal activity rates (AFA). (b) Illustration of network indices used in our analyses and their expected effects on ecosystem multifunctionality (EMF).

## Results and discussion

### Network structure affects ecosystem multifunctionality

**Fig. 2:**
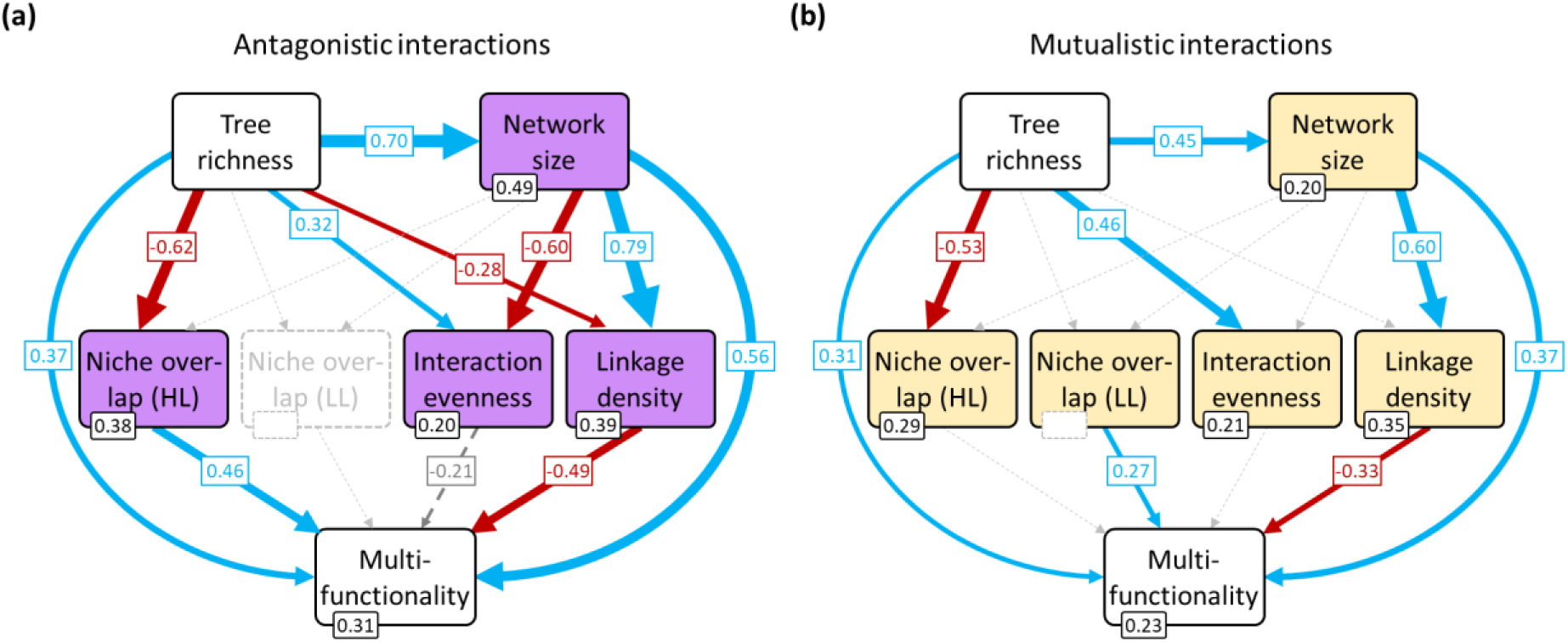
Network structure mediates the effect of tree species richness on ecosystem multifunctionality in (a) antagonistic and (b) mutualistic interaction networks. Structural equation models (SEMs) show tree species richness (i.e. the treatment in the BEF-China experiment) effects on the structure of multitrophic species interaction networks, and their joint effects on ecosystem multifunctionality. Most indices of network structure had consistently stronger direct effects on ecosystem multifunctionality than tree species richness. Positive, negative, and non-significant standardized path coefficients are indicated in blue, red, and grey, respectively. R^2^-values are displayed for each endogenous variable. HL: higher trophic level, LL: lower trophic level. Model fit measures: (a) p = 0.16, CFI = 1.00, RMSEA = 0.12, p_RMSEA_ = 0.20, SRMR = 0.05; (b) p = 0.27, CFI = 0.99, RMSEA = 0.06, p_RMSEA_ = 0.41, SRMR = 0.07. To avoid inflating the effects of antagonistic (n = 7 network types) over mutualistic (n = 4 network types) interaction networks, antagonistic and mutualistic interactions were analyzed separately (but see Supplementary Fig. 2 for overall interactions, which closely followed antagonistic interactions). Sensitivity analyses indicate that the general relationships presented here are largely invariant to individual networks (Supplementary Fig. 3a). Higher number of antagonistic networks does not affect comparability with mutualistic networks (Supplementary Fig. 3b). For an overview of model estimates, including covariances between network indices, see Supplementary Tables 4-6.

To investigate how multitrophic species interactions affect ecosystem multifunctionality, we integrated ecosystem multifunctionality, tree species richness (i.e. the experimental treatment)^42^, the number of species per interaction network (i.e. network size, corresponding to the number of interacting species), and four selected indices capturing different aspects of bipartite species interactions (i.e. abundance-weighted niche overlap of higher (HL) and lower (LL) trophic level, interaction evenness, linkage density) into two structural equation models (SEM), one for antagonistic and one for mutualistic interactions (Fig. 2). Network size and niche overlap showed positive effects, and linkage density showed negative effects on ecosystem multifunctionality. Additionally, biodiversity effects from tree species richness and network size, with the latter capturing the multi-diversity of our system (r_PEARSON_ = 0.96 between network size and mean scaled diversity across trophic levels, p < 0.001), were mediated by the other network characteristics, which was particularly pronounced for antagonistic interactions (Fig. 3b). The patterns found for antagonistic interactions (Fig. 2a) had many similarities with mutualistic interactions (Fig. 2b), leading to similar overall trends when analyzing both interaction types together (Supplementary Fig. 2), likely a result of the largely positive correlations between the structure of the different network types (Fig. 1a, left). The structure of individual network types was consistently associated with individual functions (Supplementary Table 7), with 73.8% of all possible combinations of individual network indices and functions showing significant effects of network structure. Even though our analyses integrated multiple network types that were constructed using different approaches to quantify species abundances and interaction strength (see Supplementary Table 3), and although ecosystem functions and species interactions were not perfectly temporally aligned (see Methods, Supplementary Tables 1 & 3), our general findings should be robust against these limitations. This is supported by the fact that removing interaction networks from the analyses did not change the findings (see sensitivity analyses in Supplementary Fig. 3). Thus, the multitrophic structure of species interaction networks emerges as a powerful and generalizable predictor of ecosystem multifunctionality.

Species complementarity is often considered as a main driver of biodiversity–ecosystem functioning relationships^32^ and facilitative processes have received increasing attention in recent years^46,47^. While the structure of multitrophic species interaction networks is not suited to investigate the role of facilitation, our findings indicate that niche differentiation leads to an increase in the complementarity among species, as tree species richness reduces the niche overlap of higher trophic levels and favours species with more narrow niches (i.e. specialists, low linkage density; Fig. 2). However, this did not directly translate into positive effects on ecosystem multifunctionality. While we found positive effects of narrow niches (i.e. low linkage density), a reduced niche overlap had a negative rather than the expected positive effect in antagonistic networks (Fig. 2a). In mutualistic networks, similar effects were found for niche overlap between species at the lower trophic level (Fig. 2b). So far, most of the evidence for drivers of biodiversity effects comes from studies of individual ecosystem functions^23,32,48^. While a few species can suffice to provide these functions, substantially more species are required to maintain ecosystem multifunctionality^10,49^. Hence, species may require relatively narrow niches (i.e. specialists, low linkage density) to make unique contributions to ecosystem multifunctionality. Our findings suggest that there is an additional way by which species occupying similar niches (i.e. high niche overlap) collectively contribute to ecosystem multifunctionality. High niche overlaps are traditionally associated with high competition that will eventually lead to competitive exclusion unless species differentiate their niches^33,50^. However, shared consumer species can prevent the competitive dominance of their resource species, thus enabling them to coexist despite a high niche overlap and leading to joint, more even contributions to ecosystem functioning^51,52^. The positive effects of niche overlap on ecosystem multifunctionality might therefore highlight an alternative, rarely considered complementarity mechanism that is rooted in trophic interactions and likely acts in concert with resource-use complementarity^15^.

The beneficial effects of network structure on ecosystem multifunctionality coincide with patterns associated with stability mechanisms. Specifically, niche overlaps also enable species to offset performance decreases of other species occupying similar niches (functional similarity leading to insurance effects)^35^. Especially in fluctuating environments, this can lead to asynchronous species dynamics, a form of temporal complementarity, that stabilize ecosystem functions over time^35^. Species asynchrony has been observed for plants^53^ and herbivores (Wang et al. in review) in our experiment, and similar dynamics are known from mutualists^54,55^. While the stability of ecosystem multifunctionality is not necessarily correlated with mean ecosystem multifunctionality, a positive association between stability and complementarity mechanisms such as species asynchrony can lead to synergistic effects between them^35,56,57^. Even though this cannot be tested directly, our findings therefore may also reflect a temporal dimension to species interactions that moderates biodiversity effects on ecosystem multifunctionality.

Tracing back how individual ecosystem functions determine the response of ecosystem multifunctionality to network structure reveals systematic changes of the majority of functions, with some showing contrasting trends (Supplementary Fig. 4a & 5a). The most notable responses were found for functions correlated with tree stand biomass (Supplementary Fig. 4b & 5b). At least one network index of all network types, even those not directly affecting trees (e.g. spider-prey), had significant effects on tree biomass and absolute growth rates (Supplementary Table 7), suggesting potential effects of higher-order interactions, even though the biological underpinning may differ between network types. As several ecosystem functions directly or indirectly depend on tree biomass, and tree biomass modifies the diversity at higher trophic levels^58^, we tested if tree biomass mediates effects of tree species richness on network size and ecosystem multifunctionality (Supplementary Fig. 6). While positive effects of tree species richness on tree biomass confirmed previous findings^48^, we found no significant correlation between tree biomass and network size or ecosystem multifunctionality. Hence, tree biomass cannot approximate the interplay between the multitude of processes that shape forest ecosystem functioning and are included in our study. Herbivores, for example, directly depend on the available plant biomass as a food source. However, while we found that herbivory and tree biomass both increase with network size, only tree biomass responded to niche overlap (Supplementary Fig. 4a & 5a). Previous studies could similarly show that herbivory and tree biomass are not necessarily coordinated, with processes such as predation potentially decoupling herbivory from plant productivity despite their direct association^29,33,44,59^. Our findings on tree productivity therefore also demonstrate that the effects of single bottom-up processes are limited. Instead, by utilizing network analyses to investigate drivers of biodiversity–ecosystem functioning relationships, our findings indicate that multiple processes, including species interactions, act in concert to determine ecosystem multifunctionality.

### Direct and mediated biodiversity effects

**Fig. 3:**
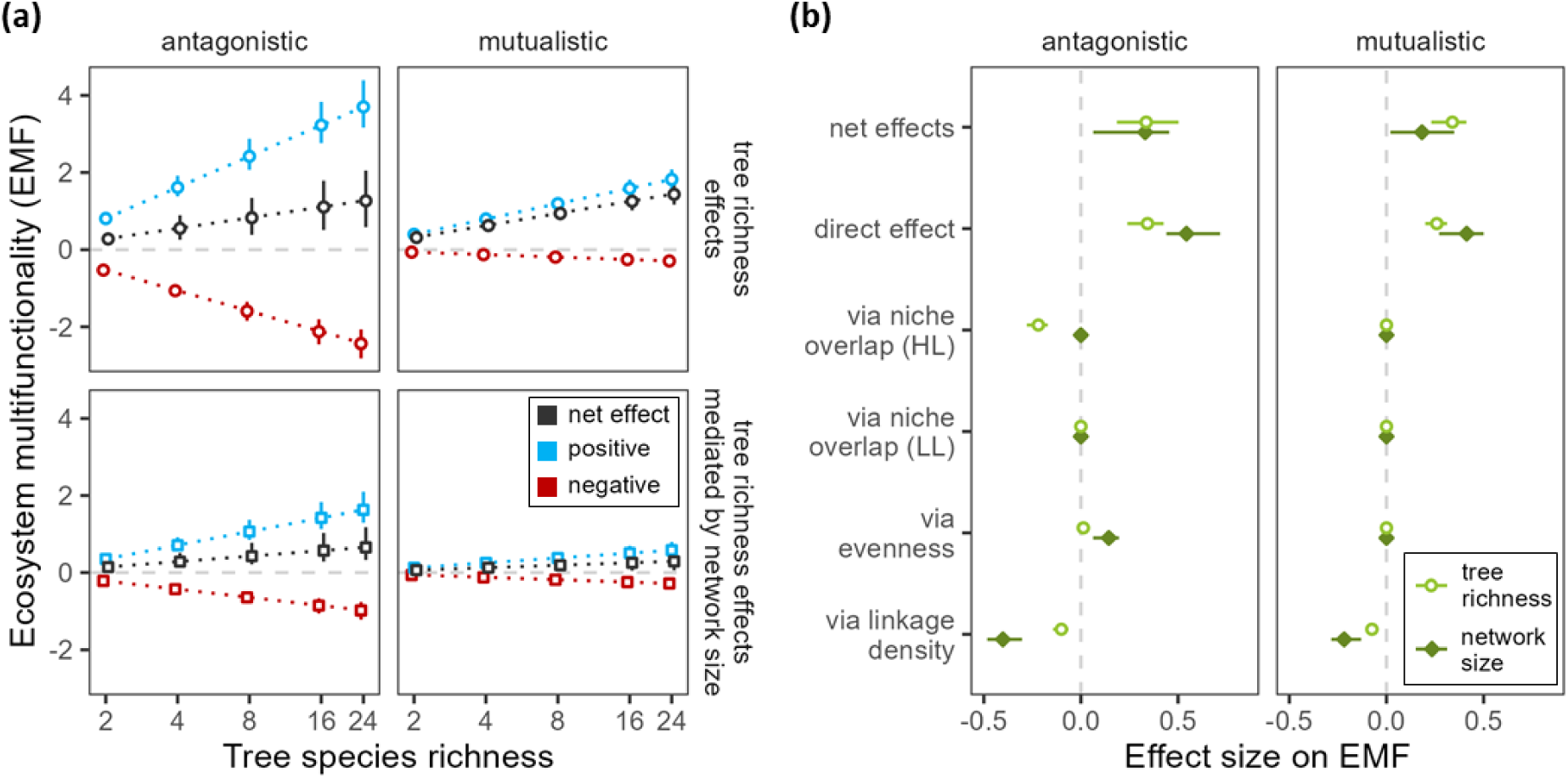
Effects of tree species richness and network size (as a measure of multitrophic diversity) on ecosystem multifunctionality (EMF). (a) Aggregated direct and indirect effects of tree species richness and network size consist of positive (blue) and negative (red) components, with net effects (black) being consistently positive. Total tree richness effects and tree richness effects mediated by network size are shown separately in rows. (b) Net, direct, and mediated effects (via niche overlap of higher (HL) and lower (LL) trophic levels, interaction evenness, linkage density) of tree species richness (light green; empty circles) and network size (dark green; filled diamonds) show that negative effects are always mediated by network structure. (a & b) Effects in antagonistic and mutualistic interaction networks are shown separately (columns). Values were calculated from path coefficients and their standard errors of the respective structural equation models (Fig. 2, Supplementary Tables 4 & 5), randomly drawn from normal distributions (N = 100). Empty squares (a), empty circles (b; tree species richness) and filled diamonds (b; network size) indicate the median. Error bars indicate interquartile ranges. A summary of the underlying values can be found in Supplementary Tables 8 & 9.

The structure of species interaction networks showed clear effects on ecosystem multifunctionality, mediating biodiversity effects of tree species richness and network size. To contextualize these effects within the total and direct effects of biodiversity on ecosystem multifunctionality, we summarized the results of our structural equation models (Fig. 2, Supplementary Tables 4 & 5) by aggregating effect sizes of tree species richness and network size (Fig. 3). Net effects of tree species richness on ecosystem multifunctionality were consistently positive, with network size mediating about half of them in antagonistic networks (Fig. 3a). In mutualistic networks, mediation via network size was smaller due to a weaker relationship with tree species richness (Fig. 2 & 3). However, for both types of interactions, network size had strong direct and indirect effects on ecosystem multifunctionality, especially in antagonistic networks (Fig. 3b). This was particularly striking given that tree species richness was manipulated experimentally and that many ecosystem functions were directly associated with trees (e.g. tree stand biomass), indicating the importance of considering species diversity across trophic levels (i.e. multi-diversity) when considering multiple ecosystem functions^19–22,37^.

In addition to direct effects on ecosystem multifunctionality, tree species richness and network size altered network structure (Fig. 2). They increased linkage density and interaction evenness, while reducing niche overlap of higher trophic levels. Together, this led to net effects of tree species richness that partitioned into positive and negative components (Fig. 3a), with the latter being mediated by changes in network structure (Fig. 3b). These findings highlight the complexity of the mechanisms underlying biodiversity effects on ecosystem multifunctionality across multiple trophic levels. The complexity is particularly evident in antagonistic networks, where tree species richness effects on multiple network characteristics were inversed by network size (Fig. 2 & 3). For linkage density, which negatively affected multifunctionality and thus indicates the importance of specialized interactions, negative effects of tree species richness increased net effects of tree species richness on ecosystem multifunctionality. However, effects mediated by network size inversed those effects, leading to net negative effects of tree species richness via linkage density. Larger networks often show higher linkage densities due to the larger number of possible interaction partners^31^, which also captures the majority of tree richness effects in our results. The fact that tree richness also reduced linkage density suggests that additional mechanisms (e.g. host dilution effects^60^) are at play. Therefore, including multiple trophic levels and their interaction structure is a crucial but often omitted step to identify the drivers influencing biodiversity effects and to capture the multitude and complexity of the underlying mechanisms^14,61^.

Compared with antagonistic interactions, mutualistic interactions only mediated tree species richness effects via network size, despite modifying network structure (Fig. 2b). This limited the indirect pathways through which tree species richness affected ecosystem multifunctionality and thus reduced the number of potential biodiversity mechanisms (Fig. 3). A lack of mediating effects of network structure may be related to the large variety of network type-specific structures displayed by mutualistic interactions (Supplementary Fig. 7). This may be a result of mutualistic network types having stronger ecological differences than antagonistic network types. For example, ant-Hemiptera interactions are rooted in an antagonistic interaction between Hemiptera and plants^62,63^, whereas interactions between symbiotic fungi and plants do not depend on other interactions but include many more species^64^. These differences suggest that responses to changes in species composition cannot be easily generalized across mutualistic networks. Accordingly, direct effects of tree species richness and network size in mutualistic networks play a more important role for generalized biodiversity effects than in antagonistic networks.

### Implications for biodiversity research and conservation

Despite some negative components, net biodiversity effects were consistently positive (Fig. 3). However, the multitude of pathways by which biodiversity altered ecosystem multifunctionality poses a challenge to predicting the consequences of diversity loss. For example, specialized species are more prone to secondary extinctions due to their narrow niches (i.e. low linkage density)^65,66^. While a loss of specialized species primarily eliminates their specific interactions, it can have secondary cascading effects by disrupting species interaction networks^66^ and redirecting energy flows^67^. Together with the benefits of specialized species for ecosystem multifunctionality, this suggests that a loss of specialized species will have disproportionally negative consequences for ecosystem functions provided by the multitrophic community. Our findings indicate that the importance of specialized interactions for ecosystem multifunctionality is consistent across types of networks and interactions. Hence, a loss of specialized species would erode ecosystem multifunctionality more than random extinction sequences. Identifying the main pathways through which species interactions mediate biodiversity effects on ecosystem multifunctionality is thus also an important tool for targeting conservation efforts. To conserve self-sustaining ecosystems, effective biodiversity conservation needs to not only focus on rare and threatened species but also target species interactions and a network structure that supports ecosystem multifunctionality^26,40,68,69^.

Our finding that ecosystem multifunctionality consistently benefits from fewer, more specialized interactions and species with similar interaction partners (i.e. low linkage density and high niche overlap) does not fully align with a niche complementarity paradigm that dominates the BEF literature^32^. Especially the positive effects of niche overlap suggest that functionally similar species can enhance ecosystem multifunctionality, indicating a potentially overlooked pathway behind positive biodiversity effects. Identifying whether this is a result of integrating multiple trophic levels and ecosystem functions, arising from a temporal niche differentiation, or caused by other mechanisms altogether should become a focus of future research. An active manipulation of network structure will be inevitable in this endeavour. While difficult under empirical settings without altering confounding factors such as species composition and diversity, theoretical models already attempted such manipulations^17,29^. To reach a complete mechanistic understanding of how network structure mediates biodiversity effects on ecosystem multifunctionality, theoretical work, however, needs to diversify the interaction and network types as well as the ecosystem functions that they entail to better match real ecosystems. Paired with identifying ways to manipulate species interactions while controlling confounding factors in experimental communities, this will enhance our ability to identify drivers of ecosystem multifunctionality.

## Methods

### Study site

All data used in this study was collected from the BEF-China tree diversity experiment, located in Xingangshan, Jiangxi Province (29°05’00’’–29°07’43’’N,117°54’19’’–117°55’53’’E) and established in 2009^42^. The experiment is characterized by a mean annual temperature of 16.7 °C and a mean annual precipitation of 1800 mm^70^. The natural vegetation in the region are highly diverse subtropical forests, comprising broadleaved evergreen and deciduous tree species. The experiment includes 40 locally occurring tree species that were planted at varying levels of diversity (1, 2, 4, 8, 16, and 24 species). In total, the experiment has 566 plots across two sites. Each plot is characterized by a defined species composition that corresponds to the levels of diversity and follows a broken-stick design^42^. In each plot, 400 tree saplings were planted in a regular 20 x 20 grid, covering a total area of 25.8 x 25.8 m². Species identities were assigned randomly to each planting position.

### Sampling

We measured a total of 34 ecosystem functions, covering 7 measures of primary production^71–75^, 19 related to nutrient cycling (nutrient resorption^76^, soil quality^77–79^, litter decomposition^73,76^, wood decomposition^80^), and 12 related to animal and fungal activity rates (4 overlapping with nutrient cycling; Fig. 1)^63,80–83^. We followed a strict definition of ecosystem functions, only selecting measures capturing ecosystem process rates^8^. All but one ecosystem function were sampled between 2014-2019, with only parasitism rates on Lepidoptera caterpillars being sampled more recently (i.e. 2021-2022). An overview of the ecosystem functions used in our analyses, including information on the sampling, can be found in Supplementary Table 1.

We assembled 11 types of species interaction networks, with 7 based on antagonistic and 4 on mutualistic interactions^63,82,84–89^. In total, 374 networks sampled between 2014-2019 directly entered our analyses. All networks are based on observed or inferred interactions between taxa of two adjacent trophic levels. We actively avoided including co-occurrence networks (e.g. by including endophytic but excluding epiphytic bacteria from the leaf bacteria data^88^). Of the 11 network types included, interactions in 6 of them are based on direct observations, for example through gut content analyses of spiders^89^, or rearing parasitoids^85^. Interactions in the remaining 5 network types are inferred, for example by assuming that symbiotic fungi sampled in the rhizosphere of a tree are interacting with the tree^87^. Details on the specific sampling and data processing protocols of the individual network types are provided in Supplementary Table 3.

### Calculating ecosystem multifunctionality

To be able to measure the simultaneous performance of multiple ecosystem functions, we utilize a multifunctionality index that captures the effective number of ecosystem functions provided by an ecosystem^43^. The approach utilizes the Hill-number framework that is also used to unify indices of species diversity^90–92^, allowing multifunctionality to take into account the number of functions, the level at which the functions are performed, and the uniformity of their performance^43^. One main advantage of effective multifunctionality is that it overcomes several limitations earlier indices have, including a loss of information when using an averaging approach, as well as a high sensitivity to data standardization, the selection of included functions, and the selection of single or multiple thresholds at which functions are considered to be provided^43^. When compared to those measures, effective multifunctionality shows strong correlations with average and threshold-based ecosystem multifunctionality, except for when the threshold is very high (i.e. 75%, where only few functions contribute to increasing multifunctionality; see Supplementary Fig. 8). Another advantage of using effective multifunctionality is that it allows to account for correlations between measures of ecosystem functions (e.g. tree growth rates and biomass) to avoid inflating the impact of shared underlying processes.

To calculate effective ecosystem multifunctionality, we multiplied the effective number of functions provided by a system with the average level at which the functions are provided. The effective number of functions was calculated using the Hill-number framework and is the number of functions that would be provided in a system where all functions are provided equally. For our analyses, we used Hill exponent q = 1 to capture inequalities between ecosystem functions in proportion to their performance, avoiding putting additional weight on low (q < 1) or high (q > 1) performing functions. We accounted for correlations between functions by incorporating correlation-based distances between functions into the calculation of the effective number of functions. For this, a distance matrix D was calculated from a correlation matrix R, where D = (1-R)/2 and values of 0 and 1 indicate perfect positive and negative correlations, respectively. By setting all values in D that fell below a threshold value τ to τ, we defined at which distance we considered two functions to be independent. We followed the recommended approach of defining τ as the average distance between functions, which aligns with a complete evaluation of all values of τ ^43,93^. In our case, this led to τ = 0.48, rendering non-correlated and negatively correlated ecosystem functions equally distinct. By avoiding to downweigh negative correlations, we are able to capture trade-offs between ecosystem functions. Once the effective number of functions was calculated, we multiplied it with the average of the standardized ecosystem functions, yielding a measure of effective ecosystem multifunctionality^43^.

To assure a meaningful comparison between ecosystem functions, ecosystem function measures EF were normalized using a min-max scaling approach, where EF’ = (EF - EF_min_) / (EF_max_ - EF_min_), with EF’ being the normalized value of a given ecosystem function, and EF, EF_min_ and EF_max_ the non-normalized, minimum and maximum values, respectively. To account for extreme outliers, EF_max_ was defined as the mean value of the three highest values. Most ecosystem functions had a true zero value where no functioning is provided. We therefore set EF_min_ to zero for all ecosystem functions other than growth rates, which could have negative values due to mortality and measurement errors.

### Capturing the structure of interaction networks

To characterize the structure of multitrophic species interactions, we constructed bipartite interaction networks^94^. Depending on the network type, interaction strength was approximated from abundances, transformed reads, or the number of observed interactions (see Supplementary Table 3). For each network, we calculated five network indices (see Supplementary Table 2 for an overview). First, network size was calculated by summing the number of species from the two adjacent trophic levels in each bipartite interaction network. Second and third, niche overlap was calculated as the abundance-weighted pairwise overlap of interactions within each trophic level, leading to two indices per network, one for each level (i.e. higher and lower), with 0 indicating no niche overlap and 1 indicating complete niche overlap. The pairwise overlap was calculated using Morisita-Horn dissimilarities^95^. Fourth, interaction evenness was calculated as the Simpson evenness of interaction strength based on the inverse of the Simpson dominance index. An interaction evenness of 0 indicates complete dominance of a single interaction and 1 indicating an equal distribution of all interactions. Finally, linkage density was calculated as the average number of species each species interacts with. We min-max scaled (between 0 and 1) network size and linkage density for comparability across networks, as the other indices are already defined between 0 and 1. We averaged network indices across the network types included in our analyses (e.g. splitting antagonistic and mutualistic interactions before averaging for the models presented in Fig. 2 & 3), an approach similarly used to capture multi-diversity^19^.

### Accounting for data heterogeneity

Since the data used in our study came from multiple sources with varying methods, we had to account for differences in sampling efforts. For ecosystem functions, this could be achieved by taking average values per plot whenever several samples were taken. For several interaction networks, it was necessary to computationally resample to standardize the sampling efforts to be the same across plots within each of the 11 network types. In such cases, we resampled each network 100 times and took average values of network indices per plot. Supplementary Table 3 gives further details on sampling efforts and resampling strategies for the individual network types.

Our analyses focused on 69 plots, where a minimum of three and an average of 5.9 types of interaction networks were sampled. To avoid inflating the inclusion criteria, we considered the three network types including soil fungi as one. We excluded monocultures to avoid trivial interaction structures of networks that include plants as a trophic level, leading to a diversity gradient ranging from 2- to 24-tree species assemblages.

To account for the heterogenous sampling of ecosystem functions and interaction networks across plots, we imputed missing data for ecosystem functions (45.18 ± 4.77 %) and network indices (50.72 ± 8.96 %) when plots were not sampled, with average error estimates of 0.00093 %. This was achieved using imputations based on predictions from random forest models. To minimize imputation noise, we ran 10,000 random forest models for each ecosystem function and network index considered in our analyses. We included topological and tree compositional information in the random forest model, which were available for all plots in our experiment. By excluding other ecosystem functions, network indices, and network types, we could avoid artificially creating associations between them. This also allowed us to use the most information available, as we were able to utilize data from plots that were excluded from our analyses but had measures of the imputed variables. Imputation error was estimated by calculating out-of-bag normalized root mean square errors. Imputations were done using the missForest package^96^ in R version 4.3.3^97^.

### Structural equation models

To analyze how the structure of species interactions mediates effects of tree species richness, i.e. the treatment in our experiment, on ecosystem multifunctionality, we used structural equation models (SEMs) that assume linear relationships between variables (using the lavaan package^98^ and R). The analyses were based on 69 sampling plots (i.e. sample size = 69) for which sampling effort was sufficient (see above). For each model, we started with a full model that included all hypothesized paths (see arrows in Fig. 2). Since we had no a-priori expectations for covariances between network indices, we investigated their importance based on modification indices and included those that indicated improvements to our model (i.e. modification index > 3.84, indicating covariances with p-values < 0.05). We then stepwise removed pathways that did not show strong influences on the overall model fit (i.e. estimates close to zero, high standard error, large p-value), making sure that model quality did not decrease. Whenever all pathways to a single variable were removed, we reassessed the covariances between network indices. When model selection led us to a candidate model, we assessed its quality using a range of model-fit measures (thresholds: p > 0.05, CFI > 0.9, RMSEA < 0.08, p_RMSEA_ > 0.05, SRMR < 0.08). Note that most fit measures in isolation are not suitable to judge on model-fit (e.g. RMSEA), which is why we also accepted models even if single fit-measures were slightly beyond their respective thresholds^99^. Before analysis, we log_2_-transformed tree species richness and checked the residuals of all included pathways. Whenever residuals were not normally distributed, we transformed the variables using a Lambert W transformation utilizing the LambertW package^100^ in R, which allows dealing with heavy-tailed and skewed variable distributions and was sufficient to normalize residuals.

## Data availability

The data underlying the analyses presented in this study will be made available after acceptance of the manuscript on a public repository.

## Acknowledgements

GA, MS, AS, HB, AE were supported by the MultiTroph Research Unit funded by the German Research Foundation (DFG, 452861007/FOR 5281). HB, BS, ST, TW acknowledge the support from the International Research Training Group TreeDì, jointly funded by the DFG (grant 319936945/GRK2324) and the University of Chinese Academy of Sciences (UCAS). We are grateful for support from the BEF-China platform, which was established with funds from the Deutsche Forschungsgemeinschaft (DFG, German Research Foundation; grant DFG FOR 891), the National Natural Science Foundation of China (NSFC 30710103907, 30930005, 31170457 and 31210103910), and the Swiss National Science Foundation (SNSF). We thank all local workers and assistants who participated in data collection. Moreover, NE acknowledges funding by the DFG (FZT 118, 202548816; Ei 862/29-1). RB acknowledges fundings by the Saxon State Ministry for Science, Culture and Tourism (SMWK) – [3-7304/35/6-2021/48880].

## Author contributions

G.A. and A.S. developed the idea, analysed the data, and led the writing. M.S. and H.B. contributed to conceptualizing the idea. All authors provided data and supported data preparation. G.A. wrote the first draft of the manuscript and all authors contributed to revisions

## References

1. Estes, J. A. et al. Trophic Downgrading of Planet Earth. Science 333, 301–306 (2011).

2. Burkle, L. A., Marlin, J. C. & Knight, T. M. Plant-Pollinator Interactions over 120 Years: Loss of Species, Co-Occurrence, and Function. Science 339, 1611–1615 (2013).

3. Shurin, J. B. et al. A cross-ecosystem comparison of the strength of trophic cascades. Ecol Lett 5, 785–791 (2002).

4. Riede, J. O. et al. Scaling of Food-Web Properties with Diversity and Complexity Across Ecosystems. in Ecological Networks, Advances in Ecological Research 42 (ed. G. Woodward), 139–170 (Academic Press Inc., 2010).

5. Buzhdygan, O. Y. et al. Biodiversity increases multitrophic energy use efficiency, flow and storage in grasslands. Nat Ecol Evol 4, 393–405 (2020).

6. Potapov, A. M. et al. Rainforest transformation reallocates energy from green to brown food webs. Nature 627, 116–122 (2024).

7. Naeem, S., Thompson, L. J., Lawler, S. P., Lawton, J. H. & Woodfin, R. M. Declining biodiversity can alter the performance of ecosystems. Nature 368, 734–737 (1994).

8. Manning, P. et al. Redefining ecosystem multifunctionality. Nat Ecol Evol 2, 427–436 (2018).

9. Barnes, A. D. et al. Energy flux: the link between multitrophic biodiversity and ecosystem functioning. Trends Ecol Evol 33, 186–197 (2018).

10. Isbell, F. et al. High plant diversity is needed to maintain ecosystem services. Nature 477, 199–202 (2011).

11. Cardinale, B. J. et al. Biodiversity loss and its impact on humanity. Nature 486, 59–67 (2012).

12. Potts, S. G. et al. Safeguarding pollinators and their values to human well-being. Nature 540, 220–229 (2016).

13. Haines-Young, R. & Potschin-Young, M. Revision of the Common International Classification for Ecosystem Services (CICES V5.1): A Policy Brief. OE 3, e27108 (2018).

14. Tilman, D., Isbell, F. & Cowles, J. M. Biodiversity and Ecosystem Functioning. Annu Rev Ecol Evol S 45, 471–493 (2014).

15. Albert, G., Gauzens, B., Loreau, M., Wang, S. & Brose, U. The hidden role of multi-trophic interactions in driving diversity–productivity relationships. Ecol Lett 25, 405–415 (2022).

16. Yu, W. et al. Systematic distributions of interaction strengths across tree interaction networks yield positive diversity–productivity relationships. Ecol Lett 27, e14338 (2024).

17. Poisot, T., Mouquet, N. & Gravel, D. Trophic complementarity drives the biodiversity-ecosystem functioning relationship in food webs. Ecol Lett 16, 853–861 (2013).

18. Giling, D. P. et al. Plant diversity alters the representation of motifs in food webs. Nat Commun 10, 1226 (2019).

19. Li, Y. et al. Plant diversity enhances ecosystem multifunctionality via multitrophic diversity. Nat Ecol Evol 8, 2037–2047 (2024).

20. Schuldt, A. et al. Biodiversity across trophic levels drives multifunctionality in highly diverse forests. Nat Commun 9, 2989 (2018).

21. Lefcheck, J. S. et al. Biodiversity enhances ecosystem multifunctionality across trophic levels and habitats. Nat Commun 6, 6936 (2015).

22. O’Connor, M. I. et al. A general biodiversity–function relationship is mediated by trophic level. Oikos 126, 18–31 (2017).

23. Fründ, J., Dormann, C. F., Holzschuh, A. & Tscharntke, T. Bee diversity effects on pollination depend on functional complementarity and niche shifts. Ecology 94, 2042–2054 (2013).

24. Wei, Z. et al. Trophic network architecture of root-associated bacterial communities determines pathogen invasion and plant health. Nat Commun 6, 8413 (2015).

25. Timóteo, S. et al. Tripartite networks show that keystone species can multitask. Funct Ecol 37, 274–286 (2023).

26. Kaiser-Bunbury, C. N. et al. Ecosystem restoration strengthens pollination network resilience and function. Nature 542, 223–227 (2017).

27. Barnosky, A. D. et al. Has the Earth’s sixth mass extinction already arrived? Nature 471, 51–57 (2011).

28. IPBES. Global Assessment Report of the Intergovernmental Science-Policy Platform on Biodiversity and Ecosystem Services. 1144 (2019).

29. Thébault, E. & Loreau, M. Food-web constraints on biodiversity-ecosystem functioning relationships. PNAS 100, 14949–14954 (2003).

30. Eschenbrenner, J. & Thébault, É. Diversity, food web structure and the temporal stability of total plant and animal biomasses. Oikos (2022) doi:10.1111/oik.08769.

31. Blüthgen, N. & Staab, M. A Critical Evaluation of Network Approaches for Studying Species Interactions. Annu Rev Ecol Evol S 55, 65–88 (2024).

32. Cardinale, B. J. et al. Impacts of plant diversity on biomass production increase through time because of species complementarity. P Natl Acad Sci USA 104, 18123–18128 (2007).

33. Amyntas, A. et al. Niche complementarity among plants and animals can alter the biodiversity–ecosystem functioning relationship. Funct Ecol 37, 2652–2665 (2023).

34. Tilman, D., Lehman, C. L. & Thomson, K. T. Plant diversity and ecosystem productivity: Theoretical considerations. P Natl Acad Sci USA 94, 1857–1861 (1997).

35. Loreau, M. & De Mazancourt, C. Biodiversity and ecosystem stability: a synthesis of underlying mechanisms. Ecol Lett 16, 106–115 (2013).

36. Schneider, F. D., Brose, U., Rall, B. C. & Guill, C. Animal diversity and ecosystem functioning in dynamic food webs. Nature Communications 7, 1–8 (2016).

37. Soliveres, S. et al. Biodiversity at multiple trophic levels is needed for ecosystem multifunctionality. Nature 536, 456–459 (2016).

38. Mello, M. A. R. et al. Keystone species in seed dispersal networks are mainly determined by dietary specialization. Oikos 124, 1031–1039 (2015).

39. Mouillot, D. et al. Rare Species Support Vulnerable Functions in High-Diversity Ecosystems. PLoS Biol 11, e1001569 (2013).

40. Tylianakis, J. M., Laliberté, E., Nielsen, A. & Bascompte, J. Conservation of species interaction networks. Biol Conserv 143, 2270–2279 (2010).

41. Thébault, E. & Fontaine, C. Stability of ecological communities and the architecture of mutualistic and trophic networks. Science 329, 853–856 (2010).

42. Bruelheide, H. et al. Designing forest biodiversity experiments: General considerations illustrated by a new large experiment in subtropical China. Methods Ecol Evol 5, 74–89 (2014).

43. Byrnes, J. E. K., Roger, F. & Bagchi, R. Understandable multifunctionality measures using Hill numbers. Oikos 2023, e09402 (2023).

44. Schmitz, O. J., Hawlena, D. & Trussell, G. C. Predator control of ecosystem nutrient dynamics. Ecology Letters 13, 1199–1209 (2010).

45. Hawlena, D., Strickland, M. S., Bradford, M. A. & Schmitz, O. J. Fear of Predation Slows Plant-Litter Decomposition. Science 336, 1434–1438 (2012).

46. Staab, M. et al. Dear neighbor: Trees with extrafloral nectaries facilitate defense and growth of adjacent undefended trees. Ecology 104, e4057 (2023).

47. Wright, A. J., Wardle, D. A., Callaway, R. & Gaxiola, A. The overlooked role of facilitation in biodiversity experiments. Trends Ecol Evol 32, 383–390 (2017).

48. Huang, Y. et al. Impacts of species richness on productivity in a large-scale subtropical forest experiment. Science 362, 80–83 (2018).

49. Hector, A. & Bagchi, R. Biodiversity and ecosystem multifunctionality. Nature 448, 188–190 (2007).

50. Chesson, P. Mechanisms of maintenance of species diversity. Annu Rev Ecol Syst 31, 343–366 (2000).

51. Brose, U. Complex food webs prevent competitive exclusion among producer species. P Roy Soc B-Biol Sci 275, 2507–2514 (2008).

52. Mortensen, B. et al. Herbivores safeguard plant diversity by reducing variability in dominance. J Ecol 106, 101–112 (2018).

53. Schnabel, F. et al. Species richness stabilizes productivity via asynchrony and drought-tolerance diversity in a large-scale tree biodiversity experiment. Sci Adv 7, eabk1643 (2021).

54. Heklau, H., Schindler, N., Eisenhauer, N., Ferlian, O. & Bruelheide, H. Temporal variation of mycorrhization rates in a tree diversity experiment. Ecol Evol 13, e10002 (2023).

55. Lázaro, A., Gómez-Martínez, C., González-Estévez, M. A. & Hidalgo, M. Portfolio effect and asynchrony as drivers of stability in plant–pollinator communities along a gradient of landscape heterogeneity. Ecography 2022, e06112 (2022).

56. Wang, S. et al. How complementarity and selection affect the relationship between ecosystem functioning and stability. Ecology 102, e03347 (2021).

57. Wagg, C. et al. Biodiversity–stability relationships strengthen over time in a long-term grassland experiment. Nat Commun 13, 7752 (2022).

58. Schuldt, A. et al. Multiple plant diversity components drive consumer communities across ecosystems. Nat Commun 10, 1460 (2019).

59. Li, Y. et al. Multitrophic arthropod diversity mediates tree diversity effects on primary productivity. Nat Ecol Evol (2023) doi:10.1038/s41559-023-02049-1.

60. Otway, S. J., Hector, A. & Lawton, J. H. Resource dilution effects on specialist insect herbivores in a grassland biodiversity experiment. Journal of Animal Ecology 74, 234–240 (2005).

61. Eisenhauer, N. et al. A multitrophic perspective on biodiversity–ecosystem functioning research. in Mechanisms underlying the relationship between biodiversity and ecosystem function, Advances in Ecological Research 61 (eds. N. Eisenhauer, D. A. Bohan, A. J. Dumbrell), 1–54 (Academic Press Inc., 2019).

62. Staab, M., Blüthgen, N. & Klein, A. M. Tree diversity alters the structure of a tri-trophic network in a biodiversity experiment. Oikos 124, 827–834 (2015).

63. Fornoff, F., Klein, A.-M., Blüthgen, N. & Staab, M. Tree diversity increases robustness of multi-trophic interactions. P Roy Soc B-Biol Sci 286, 20182399 (2019).

64. Jacoby, R., Peukert, M., Succurro, A., Koprivova, A. & Kopriva, S. The Role of Soil Microorganisms in Plant Mineral Nutrition—Current Knowledge and Future Directions. Front Plant Sci 8, 1617 (2017).

65. Schleuning, M. et al. Ecological networks are more sensitive to plant than to animal extinction under climate change. Nat Commun 7, 13965 (2016).

66. Sanders, D., Thébault, E., Kehoe, R. & Frank Van Veen, F. J. Trophic redundancy reduces vulnerability to extinction cascades. P Natl Acad Sci USA 115, 2419–2424 (2018).

67. Gilljam, D., Curtsdotter, A. & Ebenman, B. Adaptive rewiring aggravates the effects of species loss in ecosystems. Nat Commun 6, 8412 (2015).

68. Heinen, J. H., Rahbek, C. & Borregaard, M. K. Conservation of species interactions to achieve self-sustaining ecosystems. Ecography 43, 1603–1611 (2020).

69. Harvey, E., Gounand, I., Ward, C. L. & Altermatt, F. Bridging ecology and conservation: from ecological networks to ecosystem function. J Appl Ecol 54, 371–379 (2017).

70. Yang, X. et al. Establishment success in a forest biodiversity and ecosystem functioning experiment in subtropical China (BEF-China). Eur J For Res 132, 593–606 (2013).

71. Fichtner, A. et al. From competition to facilitation: how tree species respond to neighbourhood diversity. Ecol Lett 20, 892–900 (2017).

72. Germany, M. S., Bruelheide, H. & Erfmeier, A. Drivers of understorey biomass: tree species identity is more important than richness in a young forest. J Plant Ecol 14, 465–477 (2021).

73. Zhang, N. et al. Tree species richness and fungi in freshly fallen leaf litter: Unique patterns of fungal species composition and their implications for enzymatic decomposition. Soil Biol Biochem 127, 120–126 (2018).

74. Seitz, S. et al. Bryophyte-dominated biological soil crusts mitigate soil erosion in an early successional Chinese subtropical forest. Biogeosciences 14, 5775–5788 (2017).

75. Beugnon, R. et al. Abiotic and biotic drivers of tree trait effects on soil microbial biomass and soil carbon concentration. Ecol Monogr 93, e1563 (2023).

76. Deng, M. et al. Tree mycorrhizal association types control biodiversity-productivity relationship in a subtropical forest. Sci Adv 9, eadd4468 (2023).

77. Beugnon, R. et al. Tree diversity and soil chemical properties drive the linkages between soil microbial community and ecosystem functioning. ISME Comm 1, 41 (2021).

78. Scholten, T. et al. On the combined effect of soil fertility and topography on tree growth in subtropical forest ecosystems - a study from SE China. J Plant Ecol 10, 111–127 (2017).

79. Seitz, S. et al. Tree species and functional traits but not species richness affect interrill erosion processes in young subtropical forests. SOIL 2, 49–61 (2016).

80. Wu, D., Seibold, S., Pietsch, K. A., Ellwood, M. D. F. & Yu, M. Tree species richness increases spatial variation but not overall wood decomposition. Soil Biol Biochem 183, 109060 (2023).

81. Schuldt, A. et al. Herbivore and pathogen effects on tree growth are additive, but mediated by tree diversity and plant traits. Ecol Evol 7, 7462–7474 (2017).

82. Rutten, G. et al. More diverse tree communities promote foliar fungal pathogen diversity, but decrease infestation rates per tree species, in a subtropical biodiversity experiment. J Ecol 109, 2068–2080 (2021).

83. Anttonen, P. et al. Predation pressure by arthropods, birds, and rodents is interactively shaped by tree species richness, vegetation structure, and season. Front Ecol Evol 11, 1199670 (2023).

84. Wang, M. et al. Multiple components of plant diversity loss determine herbivore phylogenetic diversity in a subtropical forest experiment. J Ecol 107, 2697–2712 (2019).

85. Guo, P.-F. et al. Tree diversity promotes predatory wasps and parasitoids but not pollinator bees in a subtropical experimental forest. Basic Appl Ecol 53, 134–142 (2021).

86. Li, Y. et al. Local-scale soil nematode diversity in a subtropical forest depends on the phylogenetic and functional diversity of neighbor trees. Plant Soil 486, 441–454 (2023).

87. Singavarapu, B. et al. Tree mycorrhizal type and tree diversity shape the forest soil microbiota. Environ Microbiol 24, 4236–4255 (2022).

88. Yang, X. et al. Different assembly mechanisms of leaf epiphytic and endophytic bacterial communities underlie their higher diversity in more diverse forests. J Ecol 111, 970–981 (2023).

89. Chen, J. et al. Bottom-up and top-down effects combine to drive predator–prey interactions in a forest biodiversity experiment. Journal of Animal Ecology (2025) doi:10.1111/1365-2656.70103.

90. Hill, M. O. Diversity and Evenness: A Unifying Notation and Its Consequences. Ecology 54, 427–432 (1973).

91. Jost, L. Entropy and diversity. Oikos 113, 363–375 (2006).

92. Chao, A., Chiu, C.-H. & Jost, L. Unifying Species Diversity, Phylogenetic Diversity, Functional Diversity, and Related Similarity and Differentiation Measures Through Hill Numbers. Annu Rev Ecol Evol S 45, 297–324 (2014).

93. Chao, A. et al. An attribute-diversity approach to functional diversity, functional beta diversity, and related (dis)similarity measures. Ecol Monogr 89, e01343 (2019).

94. Dormann, C. F., Fründ, J., Blüthgen, N. & Gruber, B. Indices, Graphs and Null Models: Analyzing Bipartite Ecological Networks. Open Ecol J 2, 7–24 (2009).

95. Horn, H. S. Measurement of ‘Overlap’ in Comparative Ecological Studies. Am Nat 100, 419–424 (1966).

96. Stekhoven, D. J. & Bühlmann, P. MissForest—non-parametric missing value imputation for mixed-type data. Bioinformatics 28, 112–118 (2012).

97. R Core Team. R: A language and environment for statistical computing. R Foundation for Statistical Computing (2022).

98. Rosseel, Y. lavaan : An *R* Package for Structural Equation Modeling. J Stat Soft 48, (2012).

99. Grace, J. A ‘Weight of Evidence’ approach to evaluating structural equation models. OE 5, e50452 (2020).

100. Goerg, G. M. Lambert W random variables—a new family of generalized skewed distributions with applications to risk estimation. Ann Appl Stat 5, 2197–2230 (2011).

