## Supplementary information for "Multitrophic interaction networks mediate biodiversity effects on ecosystem multifunctionality"

**Supplementary information include:**

- Supplementary Fig. 1-8
- Supplementary Tables 1-11
- References

### Supplementary Figures

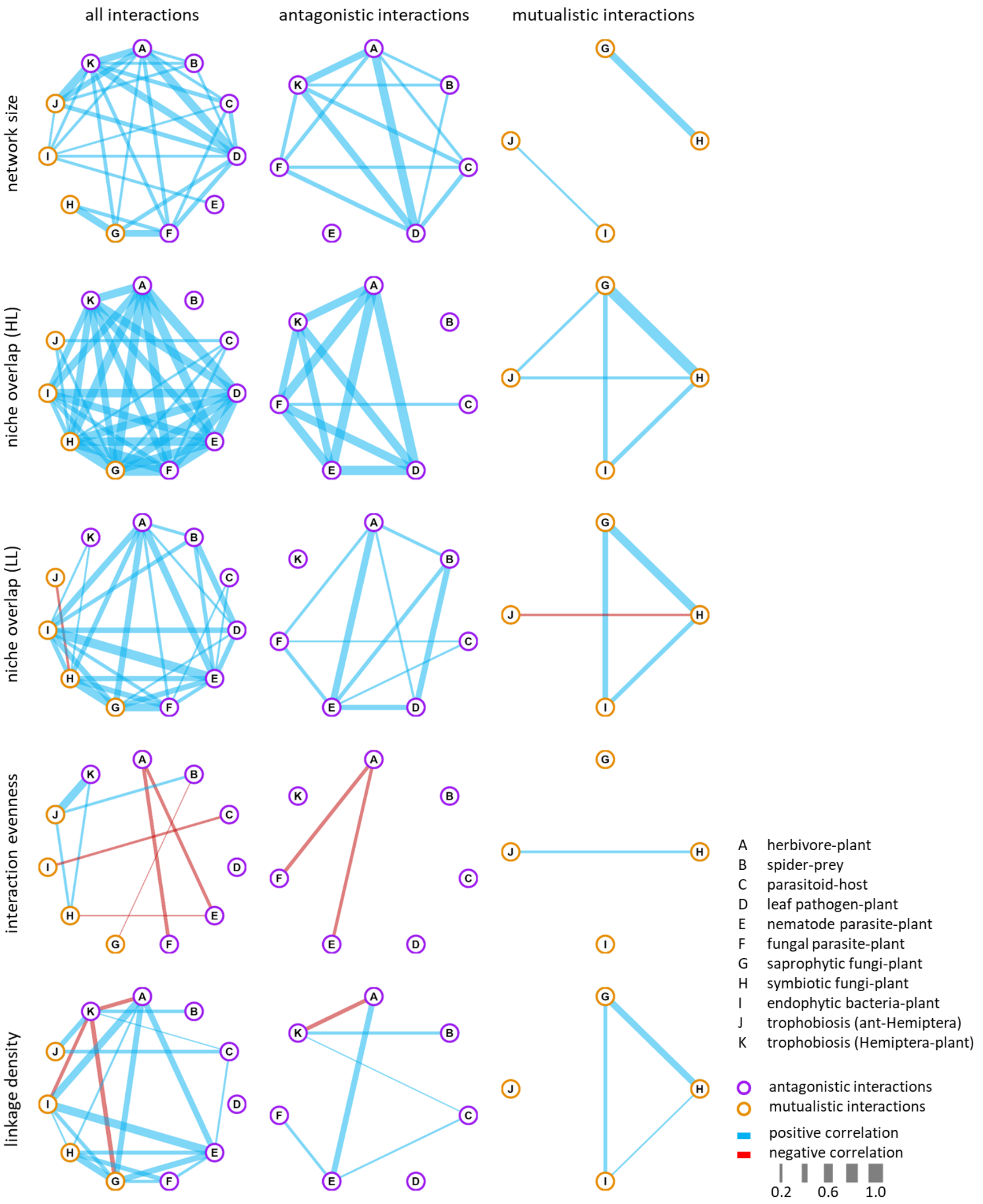


**Supplementary Fig. 1: Correlation graphs on network types.** Correlations are shown between all, only antagonistic, and only mutualistic interactions (in columns), separately for the five network indices considered (network size, niche overlap (HL), niche overlap (LL), interaction evenness, linkage density; in rows; see Supplementary Table 2 for details on their calculation). Positive correlations are indicated in blue, negative correlations in red. Linewidth scales with strength of correlation. Only significant correlations are shown (i.e. $p < 0.05$).

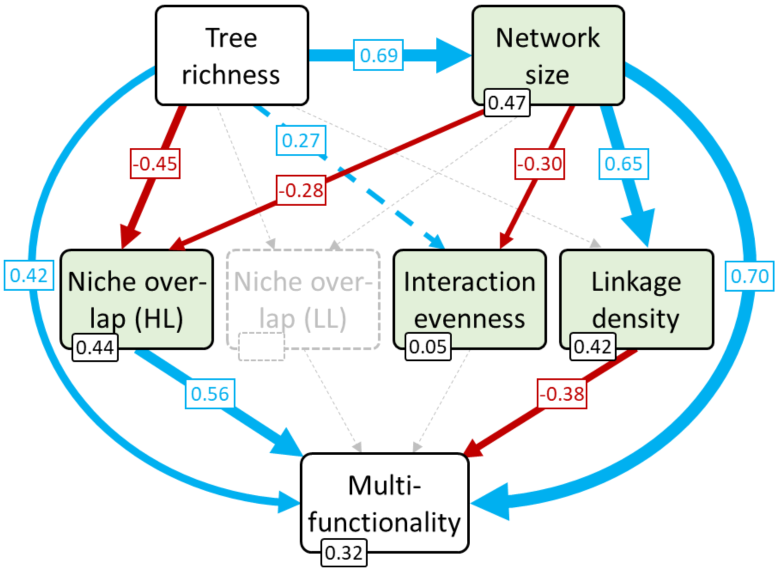


**Supplementary Fig. 2: Drivers of ecosystem multifunctionality across all interaction networks (i.e. antagonistic and mutualistic combined).** Structural equation model (SEMs) shows tree species richness (i.e. the treatment in the BEF-China experiment) effects on the structure of species interaction networks, and their joint effects on ecosystem multifunctionality. Most metrics of network structure had consistently stronger direct effects on ecosystem multifunctionality than tree species richness. Positive, negative and non-significant standardized path coefficients are indicated in blue, red, and grey, respectively. Marginally significant effects ($p < 0.1$) are indicated as red or blue dashed arrows. $R^2$-values are displayed at each endogenous variable. Model fit measures: $p = 0.39$, CFI = 1.00, RMSEA = 0.00, $p_{RMSEA} = 0.48$, SRMR = 0.02. Sensitivity analysis confirmed the robustness of the results (Supplementary Fig. 3a). For an overview of model estimates, including covariances between network indices, see Supplementary Table 6.

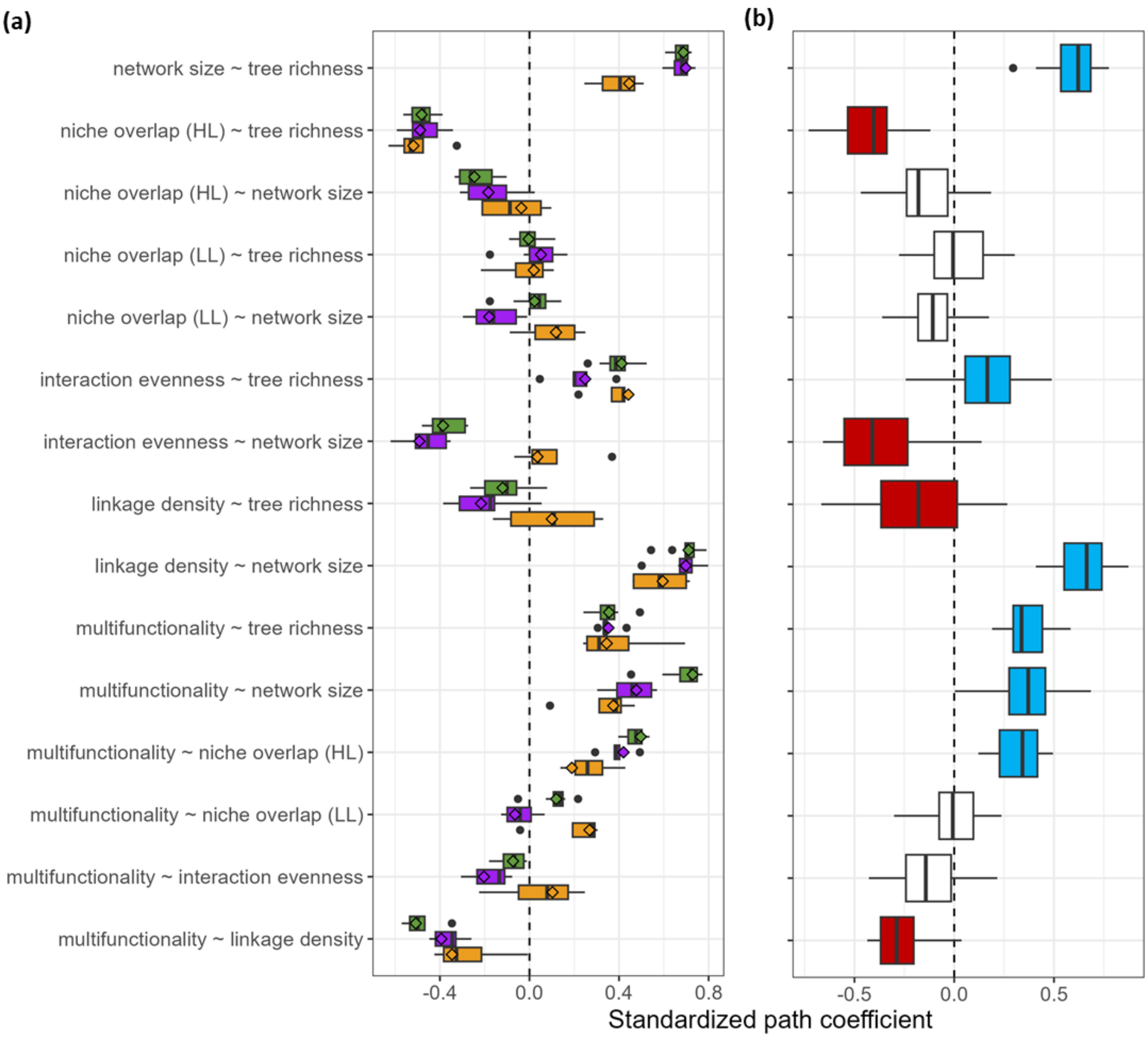


**Supplementary Fig. 3: Sensitivity analysis for results from structural equation models (SEMs).** (a) Sensitivity analysis testing the influence of individual types of species interaction networks, showing the robustness of the findings in Fig. 2 and Supplementary Fig. 2. Data from single types of interaction networks has been removed before fitting full SEMs (i.e. all pathways shown in Fig. 2 & Supplementary Fig. 2, including the non-significant ones). Standardized path coefficient of reduced (boxplot) and complete (diamonds) data are shown for each pathway for all (green; Supplementary Fig. 2, Supplementary Table 6), antagonistic (purple; Fig. 2a, Supplementary Table 4), and mutualistic interactions (yellow; Fig. 2b, Supplementary Table 5). Significant pathways are usually characterized by low spread of coefficients and/or clear tendencies to positive or negative values (e.g. network size ~ tree richness), whereas non-significant pathways show opposite patterns and often have values close to zero (e.g. niche overlap (LL) ~ tree richness). (b) Sensitivity analysis to test the influence of the higher number of types of antagonistic interaction networks (n = 7 network types) on the comparison with mutualistic interaction networks (n = 4 network types). SEMs were fitted on reduced datasets covering all 35 possible combinations of four types of antagonistic interaction networks. Effects of SEM on full dataset of antagonistic interaction networks (Fig. 2a, Supplementary Table 4) are shown in color: positive (blue), negative (red), non-significant (white). The direction of effects from reduced datasets aligns with the results of the complete dataset (Fig. 2a, Supplementary Table 4), indicating that the effects for antagonistic interaction networks are robust and therefore comparable to mutualistic interaction networks. (a & b) Full SEMs were fitted without model selection to assure comparability between them. Box in boxplots show inter-quartile range, median is displayed as thick line within box. Whiskers range to the largest or smallest value, but no further than 1.5 times the inter-quartile range away from the third and first quartile, respectively. Dots are outliers.

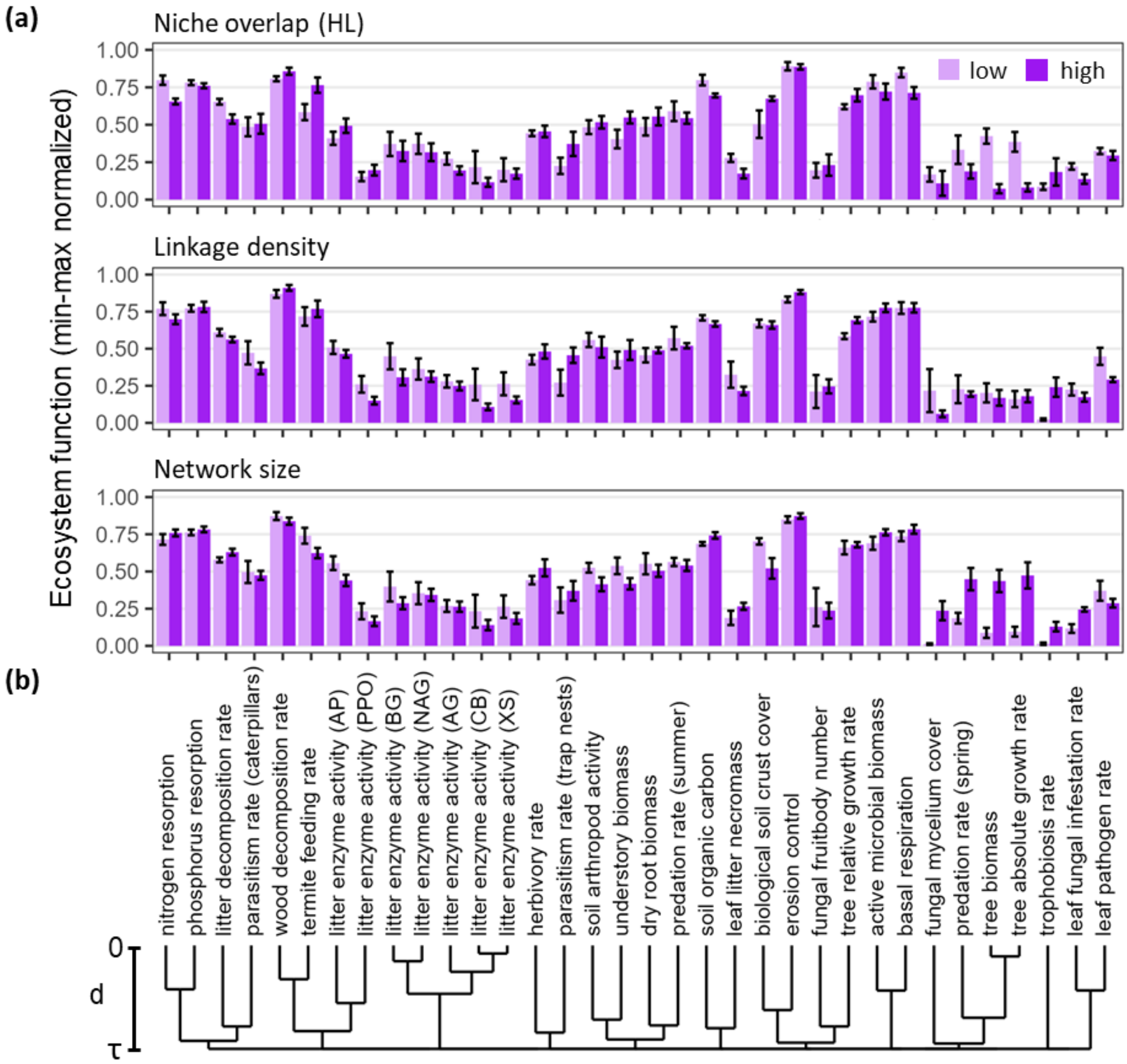


**Supplementary Fig. 4: Zooming in on how individual ecosystem functions (a) change with significant network metrics of antagonistic interactions (see Fig. 2a, Supplementary Table 4), and (b) are correlated with each other.** (a) Bar plots (mean ± SE) show values of the ten plots with the lowest (light purple) and highest (purple) niche overlap of the higher trophic level (HL; top), linkage density (middle), and network size (bottom). (b) Dendrogram of hierarchical cluster analyses based on distance matrix D = (1-R)/2, where R is the correlation matrix between ecosystem functions. Note that, compared to Fig.1a, the dendrogram was cut at τ to better represent the distances d between functions taken into account when calculating ecosystem multifunctionality (see Material and Methods for further details). The distance between functions is provided next to the dendrogram to guide interpretation.

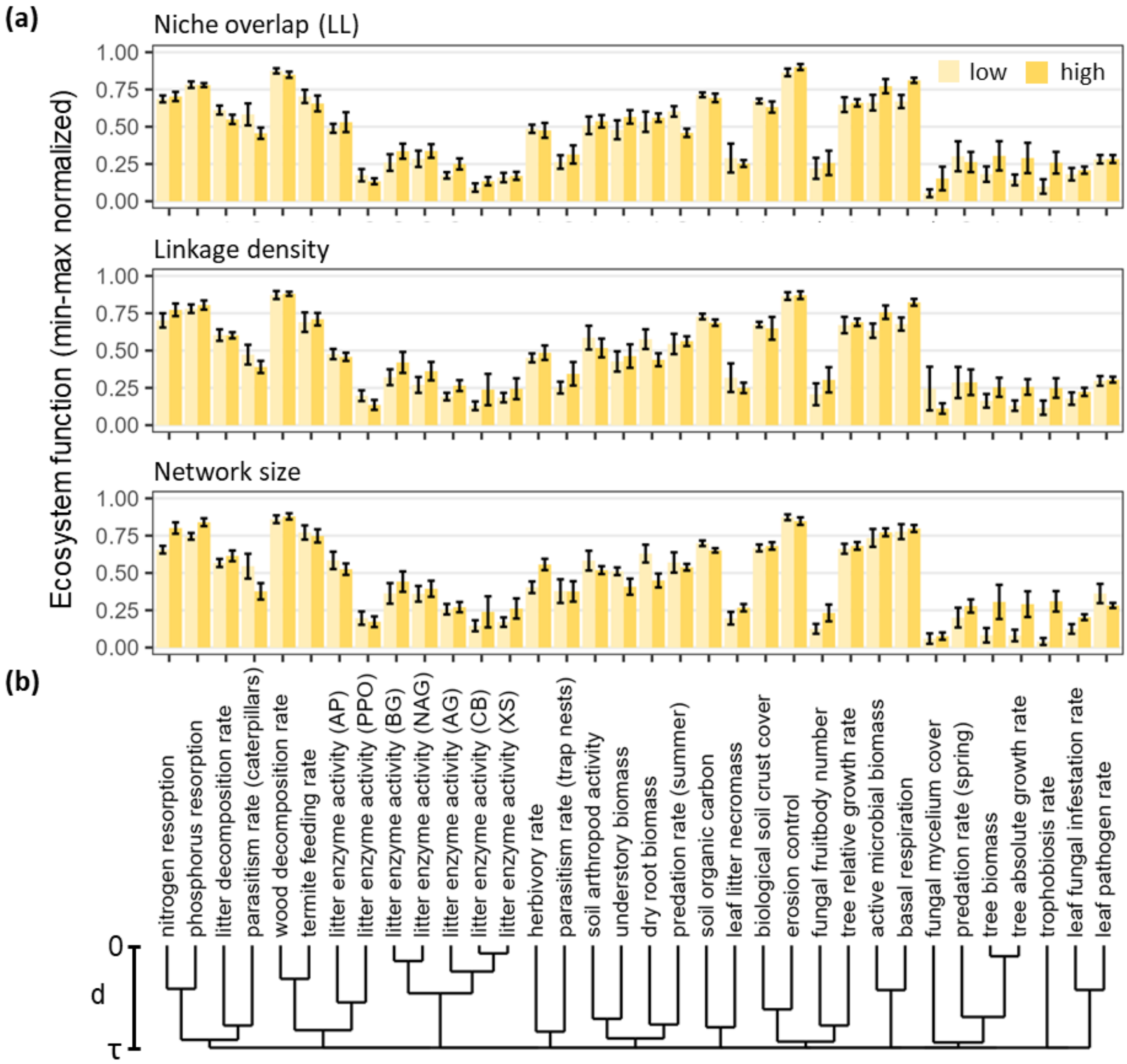


**Supplementary Fig. 5: Zooming in on how individual ecosystem functions (a) change with significant network metrics of mutualistic interactions (see Fig. 2b, Supplementary Table 5), and (b) are correlated with each other.** (a) Bar plots (mean ± SE) show values of the ten plots with the lowest (light purple) and highest (purple) niche overlap of the higher trophic level (HL; top), linkage density (middle), and network size (bottom). (b) Dendrogram of hierarchical cluster analyses based on distance matrix D = (1-R)/2, where R is the correlation matrix between ecosystem functions. Note that, compared to Fig.1a, the dendrogram was cut at τ to better represent the distances d between functions taken into account when calculating ecosystem multifunctionality (see Material and Methods for further details). The distance between functions is provided next to the dendrogram to guide interpretation.

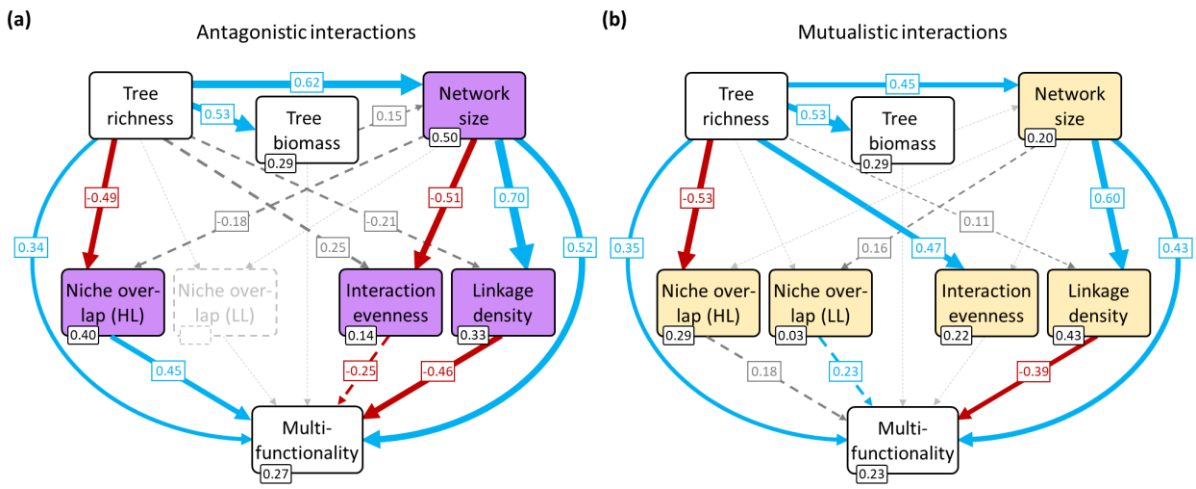


**Supplementary Fig. 6: Network structure mediates the effect of tree species richness on ecosystem multifunctionality in (a) antagonistic and (b) mutualistic interaction networks independently from tree biomass**. Structural equation models (SEMs) show tree species richness (i.e. the treatment in the BEF-China experiment) effects on the structure of multitrophic species interaction networks, and their joint effects on ecosystem multifunctionality. Compared to the SEM from Fig. 2, we explicitly tested if tree biomass mediates tree richness effects on network size and ecosystem multifunctionality. Consistent with the results presented in Fig. 2, most indices of network structure had consistently stronger direct effects on ecosystem multifunctionality than tree species richness. In contrast, tree biomass showed little to no effects. Positive, negative, and non-significant standardized path coefficients are indicated in blue, red, and grey, respectively. Marginally significant effects ($p < 0.1$) are indicated as red or blue dashed arrows. $R^2$-values are displayed for each endogenous variable. Model fit measures: (a) $p = 0.01$, CFI = 0.96, RMSEA = 0.18, $p_{RMSEA} = 0.03$, SRMR = 0.07; (b) $p = 0.20$, CFI = 0.98, RMSEA = 0.07, $p_{RMSEA} = 0.34$, SRMR = 0.07. For an overview of model estimates, including covariances between network indices, see Supplementary Tables 10 & 11.

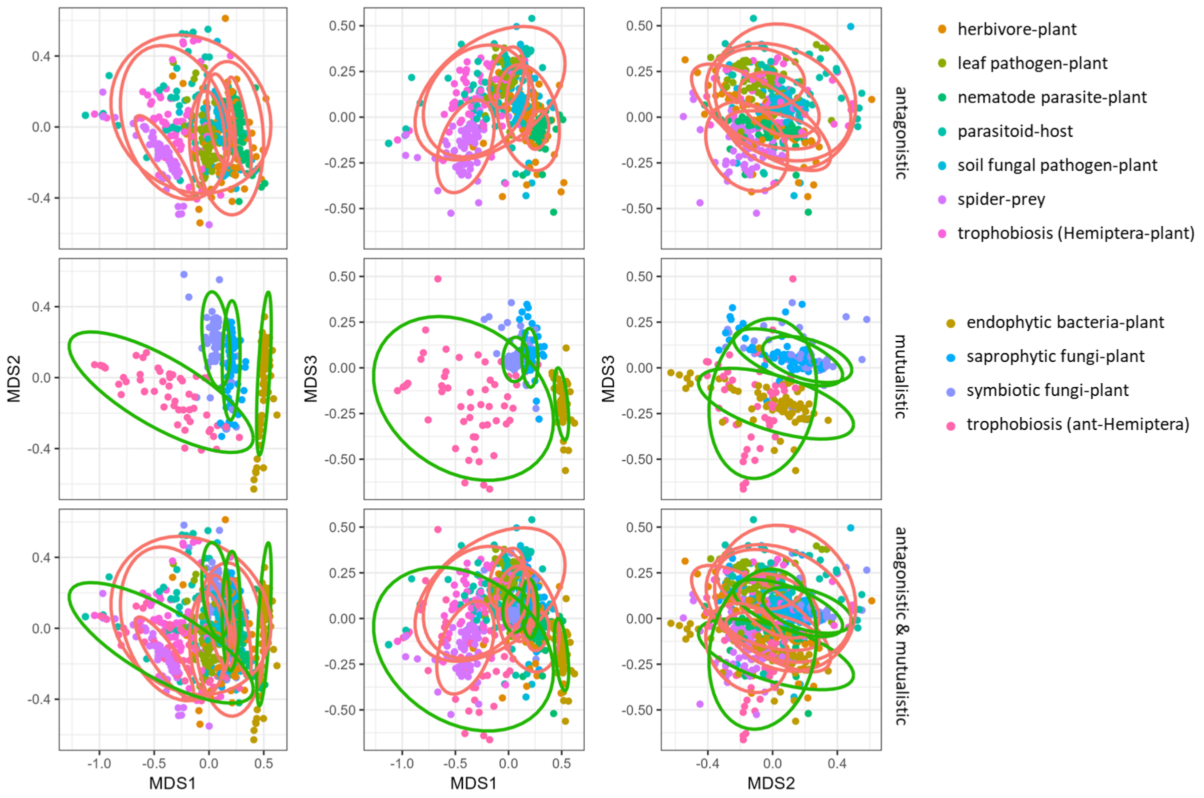


**Supplementary Fig. 7: NMDS on network characteristics (network size, niche overlap of higher and lower trophic levels, interaction evenness, linkage density).** Mutualistic interaction networks have more distinct network structures than antagonistic interaction networks. NMDS is based on Bray-Curtis dissimilarities and was calculated for all interactions together (bottom). For visual clarity, results were split for antagonistic (top) and mutualistic (middle) interactions. Stress = 0.085.

**Supplementary Fig. 8: Correlations between different ecosystem multifunctionality indices.** Indices shown are: effective ecosystem multifunctionality (eff.EMF; see Byrnes et al. 2023 and Methods for further information), as used for the analyses in this study; average ecosystem multifunctionality (mean.EMF); threshold-based ecosystem multifunctionality, at 25%, 50%, and 75% thresholds (t25, t50, t75). Pearson correlation coefficient is shown in lower triangle. Square size in upper triangle indicates absolute value of correlation coefficient (blue: positive; red: negative).

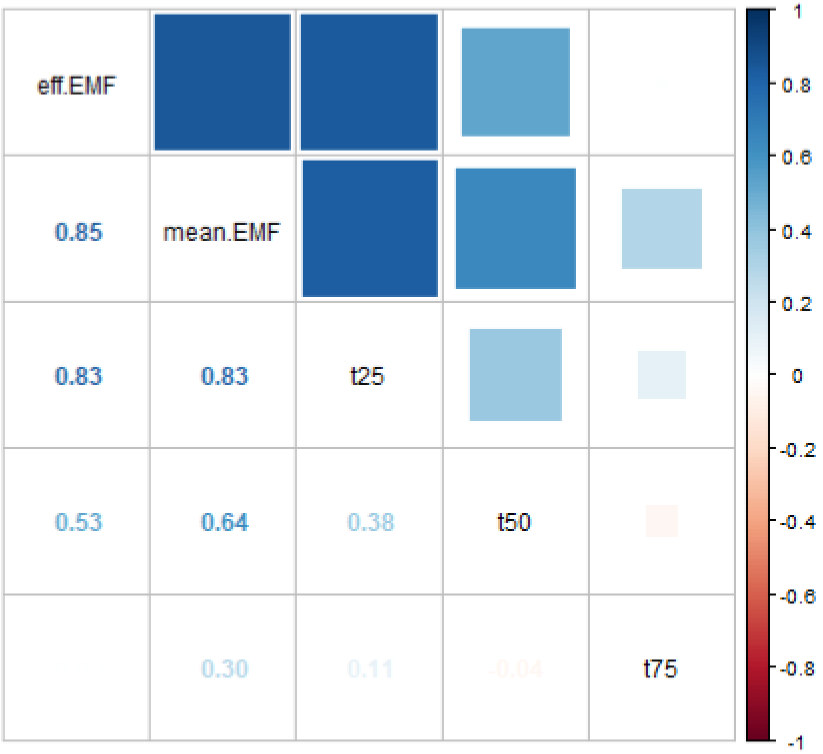

### Supplementary Tables

**Supplementary Table 1: Overview of ecosystem functions included in the analyses.** Note that all pools can be considered as rates since the start of the experiment (e.g. tree biomass) or treatment (e.g. fungal fruitbody number).

| | sampling and processing | reference |
|---|---|---|
| 1 | *tree biomass*<br>On each plot, tree height and basal area were measured from the central 36-144 tree positions depending on the diversity level. By multiplying the two measures with a form factor of 0.5, we calculated wood volume, which we subsequently multiplied with species-specific wood densities to calculate tree biomasses. We then extrapolated tree biomasses to a plot level measure, to account for differences in sampling effort.<br>year: 2016<br>plots (total / included): 495 / 69 | Fichtner et al. (2017) |
| 2 | *tree absolute growth rate*<br>Tree biomass was calculated as described above. The difference between 2016 and 2015 was recorded as absolute growth rate.<br>year: 2015-2016<br>plots (total / included): 495 / 69 | Fichtner et al. (2017) |
| 3 | *tree relative growth rate*<br>Tree biomass was calculated as described above. The difference of their log-transformed values between 2016 and 2015 was recorded as relative growth rate.<br>year: 2015-2016<br>plots (total / included): 495 / 69 | Fichtner et al. (2017) |
| 4 | *understory biomass*<br>Understory vegetation (plants < 1m in height) was harvested in August-September at three randomly positioned 0.5m x 0.5m squares at the edge of each plot, excluding trees. The weight of dried samples was used as a measure of understory biomass.<br>year: 2016<br>plots (total / included): 35 / 15 | Germany et al. (2021) |
| 5 | *leaf litter necromass*<br>Leaf litter was collected in fall using 5 litter traps per plot. Plot level litter biomass was determined after drying the mixed sample.<br>year: 2014<br>plots (total / included): 125 / 48 | Zhang et al. (2018) |
| 6 | *biological soil crust cover*<br>Biocrust cover was assessed photogrammetrically from five 0.4m x 0.4m subplots per plot and then averaged at plot level.<br>year: 2015<br>plots (total / included): 43 / 11 | Seitz et al. (2017) |

**Supplementary Table 1** – continued

| | sampling and processing | reference |
|---|---|---|
| 7 | *dry root biomass*<br>Four soil cores were taken between two neighboring trees and then mixed into one sample. The samples were sieved and root fragments were collected, air-dried, and weighted, before averaging them per plot. The number of samples varied between plots (1-11).<br>year: 2018<br>plots (total / included): 57 / 24 | Beugnon et al. (2023) |
| 8 | *nitrogen resorption*<br>Around three litter traps per plot, green and healthy leaves were collected in August of three consecutive years. Fresh litter was collected in November. After drying the samples, nitrogen concentrations were measured. The nitrogen resorption by trees was determined by comparing the nitrogen concentration of fresh litter and green leaves.<br>year: 2016-2018<br>plots (total / included): 31 / 14 | Deng et al. (2023) |
| 9 | *phosphorus resorption*<br>Phosphorus resorption was determined following the approach described for nitrogen (see above), but for phosphorus.<br>year: 2016-2018<br>plots (total / included): 31 / 14 | Deng et al. (2023) |
| 10 | *active microbial biomass*<br>Four soil cores were taken between two neighboring trees and then mixed into one sample. For each sample, active microbial biomass was measured using a substrate induced respiration method. Values were averaged to plot level. The number of samples varied between plots (1-9).<br>year: 2018<br>plots (total / included): 54 / 25 | Beugnon et al. (2021) |
| 11 | *basal respiration*<br>Soil samples were collected as described for active microbial biomass (see above). Basal respiration was measured without adding substrate, reflecting the respiration at the site.<br>year: 2018<br>plots (total / included): 54 / 25 | Beugnon et al. (2021) |
| 12 | *soil organic carbon*<br>Mixed samples of the upper 0-5 cm from 9 soil cores per plot were air-dried, grounded, and then chemically analyzed to measure soil total carbon. Since pH values were low (pH < 6.7), this is equivalent to soil organic carbon.<br>year: 2014<br>plots (total / included): 43 / 11 | Scholten et al. (2017) |

**Supplementary Table 1** – continued

| | sampling and processing | reference |
|---|---|---|
| 13 | *erosion control*<br>Soil erosion was measured on five 0.4m x 0.4m runoff plots per plot at repeated rainfall events throughout the year. It was then standardized by the amount of erosive rainfall that was locally measured. Values were extrapolated to an annual value, and averaged per plot. By sign scaling soil erosion, we received erosion control.<br>year: 2015<br>plots (total / included): 43 / 11 | Seitz et al. (2016) |
| 14 | *litter decomposition rates*<br>Litter decomposition was assessed using an in-situ decomposition experiment. Per plot, 9 litterbags were brought out and retrieved after 61, 122, and 180 days (3 replicates). After cleaning, drying and weighing the samples, decomposition rates were estimated using an exponential decomposition model on the time-series.<br>year: 2016-2017<br>plots (total / included): 31 / 15 | Deng et al. (2023) |
| 15 - 21 | *litter enzyme activity*<br>Extracellular enzyme activity was measured directly after collecting the litter as described for litter biomass (see above) using a microplate fluorometric assay. We distinguished the activity of seven enzymes (α-glucosidase (AG), β-glucosidase (BG), cellobiosidase (CB), xylosidase (XS), N-acetyl-β-glucosa-minidase (NAG) and acid phosphatase (AP), polyphenol oxidase (PPO)) as separate ecosystem functions (15-21).<br>year: 2014<br>plots (total / included): 126 / 47 | Zhang et al. (2018) |
| 22 | *wood decomposition rate*<br>Wood decomposition was determined using an in-situ experiment. Healthy branches from seven tree species representing a range of wood traits were collected. After removing the bark, samples were dried, combined into bundles, and put into 5mm mesh bags. Initial dry mass was estimated from wood volume and species-specific dry matter content. Per plot, 3 bundles were brought out at the base of 3 trees in the center of the plot. Samples were retrieved after 2 years, cleaned, dried, and weighed. Wood decomposition rates was captured as the ratio of dry mass loss to initial dry mass.<br>year: 2017<br>plots (total / included): 133 / 37 | Wu et al. (2023) |

**Supplementary Table 1** – continued

| | sampling and processing | reference |
|---|---|---|
| 24 | *fungal mycelium cover*<br>Fungal mycelium cover was assessed together with wood decomposition rates (see above). The calculation of average plot level values followed the approach for termite feeding intensity (see above).<br>year: 2017<br>plots (total / included): 133 / 37 | Wu et al. (2023) |
| 25 | *fungal fruitbody number*<br>Fungal fruitbody number was assessed together with wood decomposition rates (see above). Per sample, individual fruit bodies were counted. When fruit bodies formed conglomerations, individual basidiocarps were counted instead. Counts were averaged to plot level.<br>year: 2017<br>plots (total / included): 133 / 37 | Wu et al. (2023) |
| 26 | *soil arthropod activity*<br>Soil arthropod activity by detritivores was measured by repurposing a teabag decomposition experiment. 8 teabags were placed between two neighboring trees. Repetition varied between plots (1-11). Half of the bags were positioned under the litter layer, the rest was buried at a depth of 10 cm. Arthropod damage to teabags was assessed in 4 categories: no damage, small holes, medium holes, destroyed. To calculate a plot-level average, we transformed the 4 categories in values of 0, 10, 100, and 1000, respectively.<br>year: 2018<br>plots (total / included): 58 / 25 | Beugnon et al. (unpublished) |
| 27 | *trophobiosis rate*<br>Trees were visually assessed for the occurrence of ant-Hemiptera interactions. Trophobiosis rates are the proportion of trees where such interactions were observed out of all trees investigated per plot.<br>year: 2014<br>plots (total / included): 303 / 69 | Fornoff et al. (2019) |
| 28 | *herbivory rate*<br>Herbivore damage was assessed visually from 21 leaves per tree (7 per branch), and a total of 36-144 trees per plot, depending on plot diversity level. Damage was recorded as the relative area with visible herbivore damage, using 6 percentage classes (ranging from 0% to >75%). Average values of each class were used to calculate plot level average herbivore rates.<br>year: 2014-2015<br>plots (total / included): 280 / 68 | Schuldt et al. (2017) |

121 **Supplementary Table 1** – continued

| | sampling and processing | reference |
|---|---|---|
| 29 | *leaf pathogen rate*<br>Leaf pathogen damage was recorded following the same protocol as leaf herbivore rates (see above).<br>year: 2014-2015<br>plots (total / included): 280 / 68 | Schuldt et al. (2017) |
| 30 | *leaf fungal infestation rate*<br>Infestation rates of fungal pathogens were visually estimated on 10 randomly chosen leaves per tree as the proportion of infested area. The number of trees sampled per plot varied depending on the diversity level. Values were averaged at plot level.<br>year: 2014<br>plots (total / included): 60 / 30 | Rutten et al. (2021) |
| 31 | *predation rate (spring)*<br>Predation rates were estimated by assessing bite marks on artificial caterpillars during main activity seasons of arthropods. 6 caterpillars were placed at tree branches or the trunk of each tree. Predator attacks were recorded and caterpillars were replaced (if necessary) twice at an interval of ~1 week over a period of ~3-4 weeks. Predation rates were calculated as the proportion of caterpillars per tree that showed large bite marks. Values were averaged over all trees investigated per plot (2-22).<br>year: 2019<br>plots (total / included): 64 / 26 | Anttonen et al. (2023) |
| 32 | *predation rate (mid-summer)*<br>Measurement as for predation rates (spring; see above) but in mid-summer.<br>year: 2019<br>plots (total / included): 64 / 26 | Anttonen et al. (2023) |
| 33 | *parasitism rate (trap nests)*<br>Parasitoids were collected in trap nests as described in Supplementary Table 3. Potential hosts were raised. Parasitism rates were then calculated as the number of parasitized hosts divided by the number of potential hosts.<br>year: 2018<br>plots (total / included): 86 / 59 | Wang et al. (unpublished) |
| 34 | *parasitism rate (caterpillars)*<br>Caterpillars were collected following the protocol described for herbivore-plant interactions (s. Supplementary Table 3). All collected caterpillars were raised and the number of parasitized caterpillars was recorded. Parasitism rates were then calculated per plot as the number of parasitized caterpillars divided by the total number of caterpillars.<br>year: 2021-2022<br>plots (total / included): 52 / 28 | Wang et al. (unpublished) |

**Supplementary Table 2: Overview of network indices used in our study.** Including mathematical definitions and a brief description of ecological meaning of network indices.

| **network index** | **ecological meaning** |
|---|---|
| network size<br>$S = S_{HL} + S_{LL}$<br>$S_{HL}$ Number of species at higher trophic level<br>$S_{LL}$ Number of species at lower trophic level | The number of species interacting in a network, capturing the multitrophic species diversity. Many characteristics of species interaction networks scale with network size. |
| niche overlap (HL)<br>$NO_{HL} = \frac{2\sum_{i<j} 1 - d(x_i, x_j)}{S_{HL}(S_{HL} - 1)}$<br>$d(x_i, x_j)$ Morisita-Horn dissimilarity (Horn 1966)<br>$x_i$ Abundance of interaction partners of higher trophic level species i | Average overlap of interaction partners between species at the higher trophic level. High values indicate high niche overlaps, often associated with strong competition for resources. If values are low, species interactions are complementary. |
| niche overlap (LL)<br>$NO_{LL} = \frac{2\sum_{i<j} 1 - d(y_i, y_j)}{S_{LL}(S_{LL} - 1)}$<br>$d(y_i, y_j)$ Morisita-Horn dissimilarity (Horn 1966)<br>$y_i$ Abundance of interaction partners of lower trophic level species i | Average overlap of interaction partners between species at the lower trophic level. High values indicate high niche overlaps, which can lead to competition mediated by common consumer species (i.e. apparent competition *sensu* Holt 1977). If values are low, species interactions are complementary. |
| interaction evenness<br>$E = \frac{S}{\sum p_k^2}$ with $p_k = \frac{n_k}{\sum n_k}$<br>$n_k$ Interaction strength of interaction k | Evenness of interaction strength in a network. High values indicate that interactions are equally distributed within the network, whereas low values indicate the dominance of some interactions. |
| linkage density<br>$\bar{L}_\imath = \frac{\sum L_i}{S}$<br>$L_i$ Number of inter-actions of species i | Average number of species interactions per species, capturing how generalized (high values; species tend to interact with many species) or specialized (low values; species tend to interact with few species) species are in a network. As such, linkage density captures the average niche breadth of species in a network. |

**Supplementary Table 3: Overview of species interaction networks included in the analyses.** * Network types with inferred interactions.

| | sampling | data processing | reference |
|---|---|---|---|
| A | *herbivore-plant* * | | |
| | Lepidoptera larvae were collected by beating trees with a padded stick and capturing the falling individuals. A total of 80 trees were sampled per plot. Sampling took place three times throughout the year.<br>year: 2018<br>interaction strength: abundances<br>plots (total / included): 51 / 27<br>interaction type: antagonistic | To reduce the risk of wrongly inferring interactions, herbivore species that occurred only once per plot were removed | Wang et al. (2019) |
| B | *spider-prey* | | |
| | Spiders were sampled together with Lepidoptera larvae (see above). Gut content of spiders was then analyzed using metabarcoding methods.<br>year: 2018<br>interaction strength: number of times species was recorded in gut of spiders<br>plots (total / included): 53 / 25<br>interaction type: antagonistic | To avoid analyzing overly trivial networks, plots with less than 4 spiders were excluded | Chen et al. (2025) |
| C | *parasitoid-host* | | |
| | Hosts and their parasites were collected using trap nests positioned at the center of each sampled plot. Trap are built from multiple reed straws. Whenever reeds were sealed, they were collected and replaced. Host were directly identified. Parasitoids were identified after rearing. Trap nests were checked regularly throughout the year.<br>year: 2018<br>interaction strength: abundances<br>plots (total / included): 77 / 56<br>interaction type: antagonistic | Parasitoid abundances imputed when not recorded (~13%), using information on abundance, species, family, functional group, and sampling site | Guo et al. (2021) |
| D | *leaf pathogen-plant* | | |
| | Fungal pathogens were identified by visually assessing fungal structures of 10 randomly chosen leaves per tree using stereo- and light microscopy. The number of trees sampled per plot varied depending on the diversity level, with a minimum of 4 trees in single species assemblages.<br>year: 2014<br>interaction strength: cover<br>plots (total / included): 60 / 30<br>interaction type: antagonistic | Resampling to 4 trees per plot. One plot with 3 sampled trees was bootstrapped | Rutten et al. (2021) |

**Supplementary Table 3** - continued

| | sampling | data processing | reference |
|---|---|---|---|
| E | *nematode parasite-plant ** | | |
| | Nematodes were extracted from mixed samples of 10 soil cores taken at a distance of 10cm from a sampling tree. Nematodes were sorted into functional groups and only plant parasites were selected. The number of trees sampled per plot varied depending on the diversity level, with a minimum of 4 trees in single species assemblages.<br><br>year: 2015<br>interaction strength: abundances<br>plots (total / included): 30 / 14<br>interaction type: antagonistic | Resampling to 4 trees per plot. Abundances extrapolated from subsamples | Li et al. (2023) |
| F | *soil fungal pathogen-plant ** | | |
| | Soil fungi were sampled along the horizontal axis of focal trees using four soil cores: two at a distance of 5 cm and 25 cm from the center each. Soil cores were pooled, mixed, and sieved to yield a composite soil sample. The number of sampled trees varied between plots (2–22). From freeze-dried soil samples, microbial DNA was extracted, fungal amplicon libraries were prepared, and paired-end sequencing was performed on an Illumina MiSeq platform. Bioinformatic analysis was performed on the sequencing output to determine fungal taxa as amplicon sequence variants. Fungal taxa were annotated for potential functional groups. Only fungal pathogens were considered.<br><br>year: 2018<br>interaction strength: transformed reads<br>plots (total / included): 57 / 23<br>interaction type: antagonistic | Plots with less than 4 sampled trees were omitted. Resampling to 4 trees per plot. Fungal reads were rarefied to a sampling depth of 4,721. To avoid biases from metabarcoding (e.g. due to primer selectivity), reads were square root transformed, and rounded up before separating functional groups. | Singavarapu et al. (2022) |
| G | *saprophytic fungi-plant ** | | |
| | Sampling was done together with fungal pathogens (see above). Only saprophytic fungi were considered. Plant provides dead organic material and saprophytic fungi make nutrients available to plant they are associated with, creating a mutualistic relationship.<br><br>year: 2018<br>interaction strength: transformed reads<br>plots (total / included): 57 / 23<br>interaction type: mutualistic | Data was processed as for fungal pathogen-plant interactions | Singavarapu et al. (2022) |

**Supplementary Table 3** - continued

| | sampling | data processing | reference |
|---|---|---|---|
| H | *symbiotic fungi-plant ** | | |
| | Sampling was done together with fungal pathogens (see above). Only symbiotic fungi (e.g. mycorrhizal species) were considered.<br><br>year: 2018<br>interaction strength: transformed reads<br>plots (total / included): 57 / 23<br>interaction type: mutualistic | Data was processed as for fungal pathogen-plant interactions | Singavarapu et al. (2022) |
| I | *endophytic bacteria-plant* | | |
| | Endophytic bacteria were sampled from 10-20 shade leaves per tree and identified using metabarcoding methods. Per plot, a minimum of 8 trees were sampled. In 16- and 24-species mixtures, additional trees were sampled to contain all species in the plot.<br><br>year: 2019<br>interaction strength: transformed reads<br>plots (total / included): 29 / 15<br>interaction type: mutualistic | Metabarcoding data was rarefied to a sampling depth of 4,000, square root transformed, and rounded up. Resampling to 8 trees per plot. | Yang et al. (2023) |
| J | *ant-Hemiptera (trophobiosis)* | | |
| | Trees were visually assessed for the occurrence of ant-Hemiptera interactions. The number of trees sampled per plot varied depending on the diversity level, with a minimum of 36 trees in single species assemblages.<br><br>year: 2014<br>interaction strength: observed interactions<br>plots (total / included): 301 / 69<br>interaction type: mutualistic | Resampling to 36 trees per plot | Fornoff et al. (2019) |
| K | *Hemiptera-plant (trophobiosis)* | | |
| | Hemiptera-plant interactions were sampled together with ant-Hemiptera interactions (see above) and recorded even if ant-Hemiptera interactions were not present.<br><br>year: 2014<br>interaction strength: observed interactions<br>plots (total / included): 301 / 69<br>interaction type: antagonistic | Resampling to 36 trees per plot | Fornoff et al. (2019) |

**Supplementary Table 4: Model parameters of structural equation model (SEM) based on antagonistic interactions.** A visual representation of the SEM is shown in Fig. 2a. Model fit measures: $p = 0.16$, CFI = 1.00, RMSEA = 0.12, $p_{RMSEA} = 0.20$, SRMR = 0.05.

| | **estimate** | **std. error** | **std. estimate** | **z-value** | **p-value** |
|---|---|---|---|---|---|
| *path coefficient* | | | | | |
| network size ~ tree richness | 0.698 | 0.086 | 0.698 | 8.103 | < 0.001 |
| niche overlap (HL) ~ tree richness | -0.617 | 0.095 | -0.617 | -6.505 | < 0.001 |
| interaction evenness ~ tree richness | 0.314 | 0.135 | 0.324 | 2.321 | 0.020 |
| interaction evenness ~ network size | -0.585 | 0.123 | -0.602 | -4.763 | < 0.001 |
| linkage density ~ tree richness | -0.275 | 0.114 | -0.282 | -2.404 | 0.016 |
| linkage density ~ network size | 0.767 | 0.098 | 0.786 | 7.855 | < 0.001 |
| multifunctionality ~ tree richness | 0.375 | 0.162 | 0.372 | 2.308 | 0.021 |
| multifunctionality ~ network size | 0.566 | 0.198 | 0.562 | 2.86 | 0.004 |
| multifunctionality ~ niche overlap (HL) | 0.468 | 0.176 | 0.464 | 2.659 | 0.008 |
| multifunctionality ~ interaction evenness | -0.219 | 0.144 | -0.211 | -1.518 | 0.129 |
| multifunctionality ~ linkage density | -0.505 | 0.181 | -0.489 | -2.785 | 0.005 |
| *variance* | | | | | |
| tree richness | 0.986 | 0.168 | 1.000 | 5.874 | < 0.001 |
| network size | 0.505 | 0.086 | 0.512 | 5.874 | < 0.001 |
| multifunctionality | 0.688 | 0.117 | 0.686 | 5.874 | < 0.001 |
| niche overlap (HL) | 0.611 | 0.104 | 0.620 | 5.874 | < 0.001 |
| interaction evenness | 0.748 | 0.127 | 0.805 | 5.874 | < 0.001 |
| linkage density | 0.573 | 0.098 | 0.612 | 5.874 | < 0.001 |
| *covariance* | | | | | |
| niche overlap (HL) ~~ linkage density | 0.384 | 0.085 | 0.649 | 4.52 | < 0.001 |
| niche overlap (HL) ~~ interaction evenness | -0.369 | 0.093 | -0.546 | -3.978 | < 0.001 |
| interaction evenness ~~ linkage density | -0.356 | 0.090 | -0.544 | -3.968 | < 0.001 |

**Supplementary Table 5: Model parameters of structural equation model (SEM) based on mutualistic interactions.** A visual representation of the SEM is shown in Fig. 2b. Model fit measures: p = 0.27, CFI = 0.99, RMSEA = 0.06, $p_{RMSEA}$ = 0.41, SRMR = 0.07.

| | estimate | std. error | std. estimate | z-value | p-value |
|---|---|---|---|---|---|
| *path coefficient* | | | | | |
| network size ~ tree richness | 0.445 | 0.108 | 0.445 | 4.131 | < 0.001 |
| niche overlap (HL) ~ tree richness | -0.534 | 0.102 | -0.534 | -5.253 | < 0.001 |
| interaction evenness ~ tree richness | 0.456 | 0.101 | 0.461 | 4.506 | < 0.001 |
| linkage density ~ network size | 0.541 | 0.070 | 0.595 | 7.707 | < 0.001 |
| multifunctionality ~ tree richness | 0.310 | 0.118 | 0.309 | 2.621 | 0.009 |
| multifunctionality ~ network size | 0.374 | 0.153 | 0.372 | 2.449 | 0.014 |
| multifunctionality ~ niche overlap (LL) | 0.274 | 0.134 | 0.265 | 2.036 | 0.042 |
| multifunctionality ~ linkage density | -0.360 | 0.179 | -0.326 | -2.016 | 0.044 |
| *variance* | | | | | |
| tree richness | 0.986 | 0.168 | 1.000 | 5.874 | < 0.001 |
| network size | 0.790 | 0.135 | 0.802 | 5.874 | < 0.001 |
| multifunctionality | 0.762 | 0.130 | 0.766 | 5.874 | < 0.001 |
| niche overlap (HL) | 0.704 | 0.120 | 0.714 | 5.874 | < 0.001 |
| niche overlap (LL) | 0.930 | 0.154 | 1.000 | 6.02 | < 0.001 |
| interaction evenness | 0.758 | 0.127 | 0.787 | 5.967 | < 0.001 |
| linkage density | 0.526 | 0.090 | 0.646 | 5.874 | < 0.001 |
| *covariance* | | | | | |
| linkage density ~~ niche overlap (LL) | 0.409 | 0.095 | 0.585 | 4.31 | < 0.001 |
| niche overlap (HL) ~~ interaction evenness | -0.308 | 0.092 | -0.422 | -3.358 | < 0.001 |
| interaction evenness ~~ niche overlap (LL) | -0.201 | 0.078 | -0.240 | -2.575 | 0.010 |

**Supplementary Table 6: Model parameters of structural equation model (SEM) based on all interactions.** A visual representation of the SEM is shown in Supplementary Fig. 2. Model fit measures: p = 0.39, CFI = 1.00, RMSEA = 0.00, $p_{RMSEA}$ = 0.48, SRMR = 0.02.

| | estimate | std. error | std. estimate | z-value | p-value |
|---|---|---|---|---|---|
| *path coefficient* | | | | | |
| network size ~ tree richness | 0.688 | 0.087 | 0.688 | 7.871 | < 0.001 |
| niche overlap (HL) ~ tree richness | -0.442 | 0.105 | -0.447 | -4.21 | < 0.001 |
| niche overlap (HL) ~ network size | -0.273 | 0.115 | -0.275 | -2.381 | 0.017 |
| interaction evenness ~ tree richness | 0.253 | 0.134 | 0.266 | 1.889 | 0.059 |
| interaction evenness ~ network size | -0.288 | 0.144 | -0.304 | -1.999 | 0.046 |
| linkage density ~ network size | 0.636 | 0.090 | 0.646 | 7.037 | < 0.001 |
| multifunctionality ~ tree richness | 0.411 | 0.151 | 0.417 | 2.723 | 0.006 |
| multifunctionality ~ network size | 0.690 | 0.183 | 0.700 | 3.776 | < 0.001 |
| multifunctionality ~ niche overlap (HL) | 0.558 | 0.154 | 0.561 | 3.617 | < 0.001 |
| multifunctionality ~ linkage density | -0.382 | 0.152 | -0.381 | -2.514 | 0.012 |
| *variance* | | | | | |
| tree richness | 0.986 | 0.168 | 1.000 | 5.874 | < 0.001 |
| network size | 0.519 | 0.088 | 0.527 | 5.874 | < 0.001 |
| multifunctionality | 0.650 | 0.111 | 0.679 | 5.874 | < 0.001 |
| niche overlap (HL) | 0.538 | 0.092 | 0.556 | 5.874 | < 0.001 |
| interaction evenness | 0.841 | 0.143 | 0.948 | 5.874 | < 0.001 |
| linkage density | 0.555 | 0.095 | 0.582 | 5.874 | < 0.001 |
| *covariance* | | | | | |
| niche overlap (HL) ~~ interaction evenness | -0.362 | 0.092 | -0.538 | -3.938 | < 0.001 |
| niche overlap (HL) ~~ linkage density | 0.281 | 0.074 | 0.514 | 3.795 | < 0.001 |
| interaction evenness ~~ linkage density | -0.334 | 0.092 | -0.489 | -3.648 | < 0.001 |

**Supplementary Table 7: Effects of network structure of individual network types on individual ecosystem functions**. Based on initial structure of structural equation models (i.e. including non-significant pathways). Numbers in centre (green) count the times network indices had significant effects on each ecosystem function. Marginal numbers (red) indicate how many network types and ecosystem functions showed significant effects of network indices for each ecosystem function (rows) and network type (columns), respectively.

| | F soil fungal pathogen-plant | J trophobiosis (ant-Hemiptera) | K trophobiosis (Hemiptera-plant) | A herbivore-plant | G saprophytic fungi-plant | C parasitoid-host | H symbiotic fungi-plant | E nematode parasite-plant | D leaf pathogen-plant | B spider-prey | I endophytic bacteria-plant | # |
|---|---|---|---|---|---|---|---|---|---|---|---|---|
| basal respiration | | | | | 2 | | | | | | 1 | 2 |
| litter enzyme activity (AG) | | | | | 1 | 1 | | 1 | | 4 | 3 | 5 |
| parasitism rate (trap nests) | | | 1 | | | 1 | 2 | | 2 | 1 | 1 | 6 |
| fungal mycelium cover | | | 4 | 1 | | 1 | | 1 | | 1 | 1 | 6 |
| litter enzyme activity (XS) | | 3 | | | 1 | | 3 | 2 | | 3 | 3 | 6 |
| fungal fruitbody number | 1 | | 1 | | 1 | 1 | 1 | | | 4 | 3 | 7 |
| wood decomposition rate | | | 3 | | 2 | | 2 | 2 | 1 | 1 | 2 | 7 |
| herbivory rate | 1 | | 2 | | 1 | | 3 | 2 | 2 | | 2 | 7 |
| tree relative growth rate | | | 2 | 4 | 1 | 2 | | | 2 | 2 | 1 | 7 |
| erosion control | | | | 2 | | 4 | 1 | 2 | 5 | 1 | 1 | 7 |
| litter enzyme activity (PPO) | | 4 | | 1 | | 3 | | 2 | 2 | 3 | 3 | 7 |
| predation rate (spring) | 2 | 1 | | 1 | 2 | 1 | 2 | 2 | 3 | | | 8 |
| litter enzyme activity (AP) | 1 | 3 | 2 | | 1 | | 2 | 2 | 3 | | 1 | 8 |
| parasitism rate (caterpillars) | 1 | | | 2 | | 1 | 1 | 1 | 3 | 2 | 5 | 8 |
| leaf litter necromass | 2 | 1 | 1 | | | 2 | 3 | 3 | 4 | | 2 | 8 |
| litter enzyme activity (BG) | | 4 | 1 | | | 1 | 1 | 2 | 2 | 4 | 4 | 8 |
| litter enzyme activity (CB) | | 2 | | 2 | 3 | | 2 | 3 | 1 | 4 | 2 | 8 |
| litter enzyme activity (NAG) | | 4 | | 1 | | 2 | 1 | 3 | 2 | 3 | 3 | 8 |
| leaf fungal infestation rate | 3 | | | 1 | | 3 | 1 | 3 | 4 | 3 | 4 | 8 |
| leaf pathogen rate | | 1 | 1 | 2 | 1 | 1 | | 1 | 1 | 1 | 2 | 9 |
| dry root biomass | | | 1 | 1 | 4 | 3 | 2 | 2 | 1 | 1 | 1 | 9 |
| active microbial biomass | 2 | 1 | | 1 | 2 | | 4 | 1 | 1 | 3 | 4 | 9 |
| predation rate (summer) | | 1 | 2 | 3 | 1 | 2 | 4 | 4 | 1 | | 2 | 9 |
| biocrust cover | 2 | 3 | 3 | 2 | 1 | 3 | 2 | | 4 | 1 | | 9 |
| phosphorus resorption | | | 3 | 3 | 1 | 2 | 1 | 3 | 4 | 3 | 1 | 9 |
| termite feeding intensity | 1 | 1 | 2 | 2 | 3 | | 4 | 3 | | 4 | 1 | 9 |
| trophobiosis rate | | 2 | 1 | 1 | 1 | 2 | 1 | 2 | 2 | 2 | 3 | 10 |
| soil organic carbon | 1 | 2 | 2 | 1 | 1 | 2 | 3 | | 2 | 2 | 2 | 10 |
| nitrogen resorption | 1 | | 3 | 1 | 1 | 3 | 4 | 3 | 2 | 2 | 1 | 10 |
| soil insect activity | | 2 | 1 | 3 | 3 | 2 | 2 | 1 | 4 | 3 | 2 | 10 |
| understory biomass | 1 | | 3 | 1 | 2 | 2 | 2 | 3 | 2 | 4 | 3 | 10 |
| litter decomposition rate | 2 | | 2 | 3 | 3 | 3 | 2 | 2 | 4 | 5 | 1 | 10 |
| tree absolute growth rate | 1 | 1 | 3 | 1 | 1 | 2 | 1 | 2 | 3 | 1 | 2 | 11 |
| tree biomass | 1 | 1 | 3 | 1 | 2 | 4 | 1 | 3 | 2 | 3 | 2 | 11 |
| # | 16 | 18 | 23 | 24 | 25 | 26 | 28 | 28 | 28 | 28 | 32 | |

**Supplementary Table 8: Overview of values used in Fig. 3a.** First ($Q_1$), second ($Q_2$, median), and third ($Q_3$) quartiles are shown. Values were generated by random draws from a normal distribution, N(est., SE), based on estimates (est.) and standard errors (SE) from the fitted structural equation models (see Supplementary Tables 4 & 5).

| | 2 tree species | | | 4 tree species | | | 8 tree species | | | 16 tree species | | | 24 tree species | | |
|---|---|---|---|---|---|---|---|---|---|---|---|---|---|---|---|
| | Q1 | Q2 | Q3 | Q1 | Q2 | Q3 | Q1 | Q2 | Q3 | Q1 | Q2 | Q3 | Q1 | Q2 | Q3 |
| *antagonistic interactions - tree richness effects* | | | | | | | | | | | | | | | |
| net | 0.15 | 0.34 | 0.46 | 0.31 | 0.67 | 0.92 | 0.46 | 1.01 | 1.38 | 0.61 | 1.34 | 1.84 | 0.70 | 1.54 | 2.11 |
| pos | 0.66 | 0.81 | 0.98 | 1.32 | 1.61 | 1.96 | 1.99 | 2.42 | 2.94 | 2.65 | 3.22 | 3.92 | 3.04 | 3.70 | 4.49 |
| neg | -0.58 | -0.49 | -0.40 | -1.16 | -0.98 | -0.80 | -1.73 | -1.47 | -1.20 | -2.31 | -1.96 | -1.60 | -2.65 | -2.25 | -1.83 |
| *antagonistic interactions - network size mediated effect* | | | | | | | | | | | | | | | |
| net | 0.08 | 0.19 | 0.28 | 0.16 | 0.38 | 0.55 | 0.24 | 0.58 | 0.83 | 0.32 | 0.77 | 1.11 | 0.37 | 0.88 | 1.27 |
| pos | 0.32 | 0.40 | 0.50 | 0.63 | 0.79 | 1.00 | 0.95 | 1.19 | 1.51 | 1.27 | 1.58 | 2.01 | 1.45 | 1.82 | 2.30 |
| neg | -0.27 | -0.22 | -0.15 | -0.55 | -0.43 | -0.31 | -0.82 | -0.65 | -0.46 | -1.10 | -0.87 | -0.61 | -1.26 | -0.99 | -0.70 |
| *mutualistic interactions - tree richness effects* | | | | | | | | | | | | | | | |
| net | 0.23 | 0.33 | 0.41 | 0.46 | 0.66 | 0.81 | 0.69 | 0.99 | 1.22 | 0.91 | 1.32 | 1.63 | 1.05 | 1.51 | 1.87 |
| pos | 0.31 | 0.38 | 0.46 | 0.63 | 0.77 | 0.92 | 0.94 | 1.15 | 1.37 | 1.25 | 1.53 | 1.83 | 1.44 | 1.76 | 2.10 |
| neg | -0.10 | -0.06 | -0.03 | -0.19 | -0.12 | -0.07 | -0.29 | -0.19 | -0.10 | -0.38 | -0.25 | -0.14 | -0.44 | -0.28 | -0.16 |
| *mutualistic interactions - network size mediated effect* | | | | | | | | | | | | | | | |
| net | 0.01 | 0.07 | 0.12 | 0.02 | 0.14 | 0.23 | 0.03 | 0.22 | 0.35 | 0.04 | 0.29 | 0.46 | 0.05 | 0.33 | 0.53 |
| pos | 0.09 | 0.14 | 0.18 | 0.18 | 0.27 | 0.36 | 0.27 | 0.41 | 0.54 | 0.36 | 0.55 | 0.72 | 0.42 | 0.63 | 0.82 |
| neg | -0.09 | -0.07 | -0.04 | -0.19 | -0.14 | -0.09 | -0.28 | -0.21 | -0.13 | -0.37 | -0.27 | -0.18 | -0.43 | -0.31 | -0.20 |

**Supplementary Table 9: Overview of values used in Fig. 3b.** First ($Q_1$), second ($Q_2$, median), and third ($Q_3$) quartiles are shown. Values were generated by random draws from a normal distribution, N(est., SE), based on estimates (est.) and standard errors (SE) from the fitted structural equation models (see Supplementary Tables 4 & 5).

| | antagonistic | | | mutualistic | | |
|---|---|---|---|---|---|---|
| | Q1 | Q2 | Q3 | Q1 | Q2 | Q3 |
| *tree richness effects* | | | | | | |
| net effects | -0.01 | 0.13 | 0.31 | 0.21 | 0.26 | 0.32 |
| direct effects | 0.24 | 0.33 | 0.45 | 0.21 | 0.26 | 0.32 |
| via niche overlap (HL) | -0.28 | -0.23 | -0.17 | 0.00 | 0.00 | 0.00 |
| via niche overlap (LL) | 0.00 | 0.00 | 0.00 | 0.00 | 0.00 | 0.00 |
| via interaction evenness | -0.08 | -0.05 | -0.03 | 0.00 | 0.00 | 0.00 |
| via linkage density | 0.06 | 0.10 | 0.15 | 0.00 | 0.00 | 0.00 |
| *tree richness effects mediated by network size* | | | | | | |
| net effects | 0.18 | 0.33 | 0.52 | 0.05 | 0.21 | 0.33 |
| direct effects | 0.42 | 0.58 | 0.73 | 0.27 | 0.37 | 0.47 |
| via niche overlap (HL) | 0.00 | 0.00 | 0.00 | 0.00 | 0.00 | 0.00 |
| via niche overlap (LL) | 0.00 | 0.00 | 0.00 | 0.00 | 0.00 | 0.00 |
| via interaction evenness | 0.09 | 0.13 | 0.17 | 0.00 | 0.00 | 0.00 |
| via linkage density | -0.48 | -0.41 | -0.25 | -0.26 | -0.18 | -0.13 |

**Supplementary Table 10: Model parameters of structural equation model (SEM) based on antagonistic interactions,** including tree biomass. A visual representation of the SEM is shown in Supplementary Fig. 6a. Model fit measures: $p = 0.01$, CFI = 0.96, RMSEA = 0.18, $p_{RMSEA} = 0.03$, SRMR = 0.07.

| | **estimate** | **std. error** | **std. estimate** | **z-value** | **p-value** |
|---|---|---|---|---|---|
| *path coefficient* | | | | | |
| tree biomass ~ tree richness | 0.453 | 0.086 | 0.533 | 5.239 | < 0.001 |
| network size ~ tree richness | 0.617 | 0.1 | 0.617 | 6.154 | < 0.001 |
| network size ~ tree biomass | 0.181 | 0.118 | 0.153 | 1.529 | 0.126 |
| niche overlap (HL) ~ tree richness | -0.488 | 0.131 | -0.488 | -3.741 | < 0.001 |
| niche overlap (HL) ~ network size | -0.183 | 0.131 | -0.183 | -1.405 | 0.160 |
| interaction evenness ~ tree richness | 0.237 | 0.146 | 0.253 | 1.625 | 0.104 |
| interaction evenness ~ network size | -0.474 | 0.146 | -0.506 | -3.249 | 0.001 |
| linkage density ~ tree richness | -0.206 | 0.138 | -0.206 | -1.496 | 0.135 |
| linkage density ~ network size | 0.697 | 0.138 | 0.698 | 5.061 | < 0.001 |
| multifunctionality ~ tree richness | 0.333 | 0.157 | 0.336 | 2.118 | 0.034 |
| multifunctionality ~ network size | 0.512 | 0.200 | 0.517 | 2.553 | 0.011 |
| multifunctionality ~ niche overlap (HL) | 0.447 | 0.178 | 0.451 | 2.503 | 0.012 |
| multifunctionality ~ interaction evenness | -0.266 | 0.146 | -0.252 | -1.821 | 0.069 |
| multifunctionality ~ linkage density | -0.457 | 0.169 | -0.461 | -2.696 | 0.007 |
| *variance* | | | | | |
| tree richness | 0.986 | 0.168 | 1.000 | 5.874 | < 0.001 |
| tree biomass | 0.508 | 0.086 | 0.715 | 5.874 | < 0.001 |
| network size | 0.488 | 0.083 | 0.496 | 5.874 | < 0.001 |
| multifunctionality | 0.703 | 0.120 | 0.728 | 5.874 | < 0.001 |
| niche overlap (HL) | 0.594 | 0.101 | 0.603 | 5.874 | < 0.001 |
| interaction evenness | 0.742 | 0.126 | 0.859 | 5.874 | < 0.001 |
| linkage density | 0.660 | 0.112 | 0.672 | 5.874 | < 0.001 |
| *covariance* | | | | | |
| niche overlap (HL) ~~ linkage density | 0.400 | 0.089 | 0.639 | 4.472 | < 0.001 |
| interaction evenness ~~ linkage density | -0.379 | 0.096 | -0.542 | -3.958 | < 0.001 |
| niche overlap (HL) ~~ interaction evenness | -0.358 | 0.091 | -0.540 | -3.947 | < 0.001 |

**Supplementary Table 11: Model parameters of structural equation model (SEM) based on mutualistic interactions,** including tree biomass. A visual representation of the SEM is shown in Supplementary Fig. 6b. Model fit measures: p = 0.20, CFI = 0.98, RMSEA = 0.07, $p_{RMSEA}$ = 0.34, SRMR = 0.07.

| | **estimate** | **std. error** | **std. estimate** | **z-value** | **p-value** |
|---|---|---|---|---|---|
| *path coefficient* | | | | | |
| tree biomass ~ tree richness | 0.453 | 0.086 | 0.533 | 5.239 | < 0.001 |
| network size ~ tree richness | 0.445 | 0.108 | 0.445 | 4.131 | < 0.001 |
| niche overlap (HL) ~ tree richness | -0.534 | 0.102 | -0.534 | -5.253 | < 0.001 |
| niche overlap (LL) ~ network size | 0.154 | 0.112 | 0.159 | 1.378 | 0.168 |
| interaction evenness ~ tree richness | 0.462 | 0.103 | 0.467 | 4.498 | < 0.001 |
| linkage density ~ tree richness | 0.101 | 0.078 | 0.106 | 1.282 | 0.200 |
| linkage density ~ network size | 0.570 | 0.093 | 0.599 | 6.094 | < 0.001 |
| multifunctionality ~ tree richness | 0.343 | 0.136 | 0.345 | 2.516 | 0.012 |
| multifunctionality ~ network size | 0.431 | 0.149 | 0.433 | 2.886 | 0.004 |
| multifunctionality ~ niche overlap (HL) | 0.181 | 0.125 | 0.182 | 1.454 | 0.146 |
| multifunctionality ~ niche overlap (LL) | 0.239 | 0.135 | 0.234 | 1.775 | 0.076 |
| multifunctionality ~ linkage density | -0.408 | 0.180 | -0.390 | -2.271 | 0.023 |
| *variance* | | | | | |
| tree richness | 0.986 | 0.168 | 1.000 | 5.874 | < 0.001 |
| tree biomass | 0.508 | 0.086 | 0.715 | 5.874 | < 0.001 |
| network size | 0.790 | 0.135 | 0.802 | 5.874 | < 0.001 |
| multifunctionality | 0.754 | 0.128 | 0.773 | 5.874 | < 0.001 |
| niche overlap (HL) | 0.704 | 0.120 | 0.714 | 5.874 | < 0.001 |
| niche overlap (LL) | 0.907 | 0.150 | 0.975 | 6.025 | < 0.001 |
| interaction evenness | 0.756 | 0.127 | 0.782 | 5.971 | < 0.001 |
| linkage density | 0.511 | 0.087 | 0.574 | 5.874 | < 0.001 |
| *covariance* | | | | | |
| niche overlap (LL) ~~ linkage density | 0.395 | 0.092 | 0.581 | 4.293 | < 0.001 |
| niche overlap (HL) ~~ interaction evenness | -0.306 | 0.091 | -0.420 | -3.347 | 0.001 |
| niche overlap (LL) ~~ interaction evenness | -0.204 | 0.078 | -0.246 | -2.626 | 0.009 |

### References


Anttonen, P. *et al.* Predation pressure by arthropods, birds, and rodents is interactively shaped by tree species richness, vegetation structure, and season. *Front Ecol Evol* **11**, 1199670 (2023).

Beugnon, R. *et al.* Tree diversity and soil chemical properties drive the linkages between soil microbial community and ecosystem functioning. *ISME Comm* **1**, 41 (2021).

Beugnon, R. *et al.* Abiotic and biotic drivers of tree trait effects on soil microbial biomass and soil carbon concentration. *Ecol Monogr* **93**, e1563 (2023).

Byrnes, J. E. K., Roger, F. & Bagchi, R. Understandable multifunctionality measures using Hill numbers. *Oikos* **2023**, e09402 (2023).

Chen, J. *et al.* Bottom-up and top-down effects combine to drive predator–prey interactions in a forest biodiversity experiment. *Journal of Animal Ecology* (2025) doi:10.1111/1365-2656.70103.

Deng, M. *et al.* Tree mycorrhizal association types control biodiversity-productivity relationship in a subtropical forest. *Sci Adv* **9**, eadd4468 (2023).

Fichtner, A. *et al.* From competition to facilitation: how tree species respond to neighbourhood diversity. *Ecol Lett* **20**, 892–900 (2017).

Fornoff, F., Klein, A.-M., Blüthgen, N. & Staab, M. Tree diversity increases robustness of multi-trophic interactions. *P Roy Soc B-Biol Sci* **286**, 20182399 (2019).

Germany, M. S., Bruelheide, H. & Erfmeier, A. Drivers of understorey biomass: tree species identity is more important than richness in a young forest. *J Plant Ecol* **14**, 465–477 (2021).

Guo, P.-F. *et al.* Tree diversity promotes predatory wasps and parasitoids but not pollinator bees in a subtropical experimental forest. *Basic Appl Ecol* **53**, 134–142 (2021).

Holt, R. D. Predation, Apparent Competition, and the Structure of Prey Communities. *Theor Popul Biol* **12**, 197–229 (1977).

Horn, H. S. Measurement of ‘Overlap’ in Comparative Ecological Studies. *Am Nat* **100**, 419–424 (1966).

Li, Y. *et al.* Local-scale soil nematode diversity in a subtropical forest depends on the phylogenetic and functional diversity of neighbor trees. *Plant Soil* **486**, 441–454 (2023).

Rutten, G. *et al.* More diverse tree communities promote foliar fungal pathogen diversity, but decrease infestation rates per tree species, in a subtropical biodiversity experiment. *J Ecol* **109**, 2068–2080 (2021).

Scholten, T. *et al.* On the combined effect of soil fertility and topography on tree growth in subtropical forest ecosystems’a study from SE China. *J Plant Ecol* **10**, 111–127 (2017).

Schuldt, A. *et al.* Herbivore and pathogen effects on tree growth are additive, but mediated by tree diversity and plant traits. *Ecol Evol* **7**, 7462–7474 (2017).

Seitz, S. *et al.* Tree species and functional traits but not species richness affect interrill erosion processes in young subtropical forests. *SOIL* **2**, 49–61 (2016).

Seitz, S. *et al.* Bryophyte-dominated biological soil crusts mitigate soil erosion in an early successional Chinese subtropical forest. *Biogeosciences* **14**, 5775–5788 (2017).

Singavarapu, B. *et al.* Tree mycorrhizal type and tree diversity shape the forest soil microbiota. *Environ Microbiol* **24**, 4236–4255 (2022).

Wang, M. *et al.* Multiple components of plant diversity loss determine herbivore phylogenetic diversity in a subtropical forest experiment. *J Ecol* **107**, 2697–2712 (2019).

Wu, D., Seibold, S., Pietsch, K. A., Ellwood, M. D. F. & Yu, M. Tree species richness increases spatial variation but not overall wood decomposition. *Soil Biol Biochem* **183**, 109060 (2023).

Yang, X. *et al.* Different assembly mechanisms of leaf epiphytic and endophytic bacterial communities underlie their higher diversity in more diverse forests. *J Ecol* **111**, 970–981 (2023).

Zhang, N. *et al.* Tree species richness and fungi in freshly fallen leaf litter: Unique patterns of fungal species composition and their implications for enzymatic decomposition. *Soil Biol Biochem* **127**, 120–126 (2018).
